# Basis of Specificity for a Conserved and Promiscuous Chromatin Remodeling Protein

**DOI:** 10.1101/2020.05.25.115584

**Authors:** Drake A. Donovan, Johnathan G. Crandall, Vi N. Truong, Abigail L. Vaaler, Thomas B. Bailey, Devin Dinwiddie, Laura E. McKnight, Jeffrey N. McKnight

## Abstract

Eukaryotic genomes are organized dynamically through the repositioning of nucleosomes. Isw2 is an enzyme that has been previously defined as a genome-wide, non-specific nucleosome spacing factor. Here, we show that Isw2 instead acts as an obligately targeted nucleosome remodeler *in vivo* through physical interactions with sequence-specific factors. We demonstrate that Isw2-recruiting factors use small and previously uncharacterized epitopes, which direct Isw2 activity through highly conserved acidic residues in the Isw2 accessory protein Itc1. This interaction orients Isw2 on target nucleosomes, allowing for precise nucleosome positioning at targeted loci. Finally, we show that these critical acidic residues have been lost in the *Drosophila* lineage, potentially explaining the inconsistently characterized function of Isw2-like proteins. Altogether, these data suggest an “interacting barrier model” where Isw2 interacts with a sequence-specific factor to accurately and reproducibly position a single, targeted nucleosome to define the precise border of phased chromatin arrays.

## Introduction

Chromatin consists of the nucleic acids and proteins that make up the functional genome of all eukaryotic organisms. The most basic regulatory and structural unit of chromatin is the nucleosome. Each nucleosome is defined as an octamer of histone proteins, which is wrapped by approximately 147 base pairs of genomic DNA^1,2^. The specific positioning of nucleosomes on the underlying DNA can have significant effects on downstream processes, such as promoter accessibility and molecular recruitment, which ultimately serve to alter gene expression^3^. Despite decades of research, the mechanisms leading to precise nucleosome locations in cells are still being defined.

Nucleosome positioning is dynamically established by a group of enzymes known as ATP-dependent chromatin remodeling proteins^4^ (ChRPs). Extensive biochemical and structural characterization has been performed on this group of proteins from various families^5^. The CHD and ISWI families of ChRPs have been characterized as nonspecific nucleosome sliding and spacing factors *in vitro*^6–12^. In yeast, flies, and mammals ChRPs generate evenly spaced nucleosome arrays at transcription start sites (TSSs) and organize genomic chromatin at other defined boundaries^12–20^. However, relatively little is known about the *in vivo* biological regulation of these spacing factors, and it is not understood how they can accurately and reproducibly position nucleosomes throughout the genome in different cellular contexts.

A widely accepted model is that ChRPs pack nucleosome arrays against a non-interacting barrier, such as an unrelated DNA-binding protein or another nucleosome^16,20,21^. In this way, general regulatory factors (GRFs) could establish chromatin landscapes with differing nucleosome arrays in response to changes in the cellular environment. In support of this model, nucleosome arrays near GRFs and other DNA binding elements appear to be phased relative to the binding motifs of the sequence-specific DNA binding factors in cells and in biochemically-reconstituted cell-free systems^16,18,22^. This model suggests that boundaries of nucleosome arrays are determined by the binding of barrier factors. Implicit in this barrier model are the assumptions that ChRPs act as nonspecific nucleosome spacing machines throughout the genome and that specific ChRP and GRF interactions are not required to establish nucleosome positions. While this model provides a good explanation for how phased nucleosome arrays can be established throughout the genome by a combination of DNA binding factors and nonspecific chromatin remodeling factors, the fundamental assumptions of the barrier model have not been thoroughly tested.

It has been shown through genetic and recent biochemical experiments that members of the ISWI family of ChRPs functionally interact with transcription factors *in vivo*^16,23–26^. One of the most well-defined interacting partners of ISWI proteins is the meiotic repressor Ume6, which is found in yeasts. It has been previously demonstrated that Ume6 and Isw2, an ISWI-containing ChRP complex in *S. cerevisiae* (homologous to the ACF complex in humans and flies), share genetic targets of repression and likely interact physically^24^. While interactions with sequence-specific DNA binding proteins can potentially determine precise nucleosome targeting and final nucleosome positions^27–29^, the mechanisms through which physical interactions between Isw2 and any genomic recruitment factor like Ume6 influence nucleosome positioning activity in cells has not been defined. For example, it is not known how these physical interactions occur or what role they play in the biochemical outcomes of chromatin remodeling reactions and the resulting downstream biological outputs.

In this work, we have successfully identified the mechanism of interaction between Isw2 and Ume6 in *Saccharomyces cerevisiae*. By taking a protein dissection approach combined with genome-wide nucleosome profiling, we have identified a previously uncharacterized helical domain in Ume6 that allows for Isw2 binding, specific genomic recruitment, and precise nucleosome positioning outcomes. We further demonstrate that conserved attributes of this helical domain are observed in the cell cycle regulator Swi6, which we have identified as a new Isw2-recruitment adapter protein that allows for specific nucleosome positioning at Mbp1/Swi6 (MBF) and Swi4/Swi6 (SBF) targets. We have also determined that the transcription factor-interacting interface of Isw2/ACF-like remodeling complexes contains a few key and highly conserved residues within the WAC domain. Finally, we show that these residues, which are essential for directional, sequence-specific remodeling, were lost in the evolution of the *Drosophila* lineage, where extensive biochemical, genetic and genomic characterization has been performed on the *ITC1* ortholog ACF.

## Results

### Isw2 Activity in Cells is Inconsistent with Known Biochemistry and the Barrier Model for Nucleosome Packing

We wished to understand how the conserved Isw2 protein complex in yeast behaves genome-wide and at specific promoter nucleosomes at target sites. Yeast Isw2 has been characterized extensively in biochemical assays, which all suggest that it has nonspecific DNA binding, ATP hydrolysis, nucleosome sliding, mononucleosome centering and nucleosome spacing activities^6,9–11,30–35^. These nonspecific nucleosome mobilizing activities suggest that the Isw2 protein should be able to organize nucleosome arrays against a barrier across the genome in yeast cells, since 1) it is estimated that there are enough Isw2 molecules for every 10-20 nucleosomes in the genome^23^, 2) *D. melanogaster* ACF, an Isw2 ortholog, can organize nucleosomes into evenly-spaced arrays^18^, and 3) other nonspecific and related nucleosome spacing factors can globally space nucleosomes across the genome in yeast and other organisms^12,17,19,20^. To first determine how Isw2 positions nucleosomes in *S. cerevisiae*, we examined nucleosome positioning activity in an *isw1/chd1* deletion background to remove known and potentially overlapping global spacing factors and highlight “isolated positioning activity” by Isw2. When examining the positioning of nucleosomes with and without Isw2 at all yeast pre-initiation complex sites (PICs), it is evident that Isw2 activity is specialized at only a subset of target sites (Figure 1A). As seen previously^19,36^, no global nucleosome spacing or organizing activity is detected by Isw2 alone (Figure S1A). Close inspection of Isw2-targeted PICs suggests that Isw2 can only organize a single PIC-proximal nucleosome, while subsequent nucleosomes become more poorly phased as the distance from the initially positioned nucleosome increases (Figure 1A, Figure S1B). Importantly, the PICs that display specific Isw2-directed activity are bound by Isw2, while those lacking any detectable nucleosome organization by Isw2 are unbound (Figure S1C).

**Figure 1.**
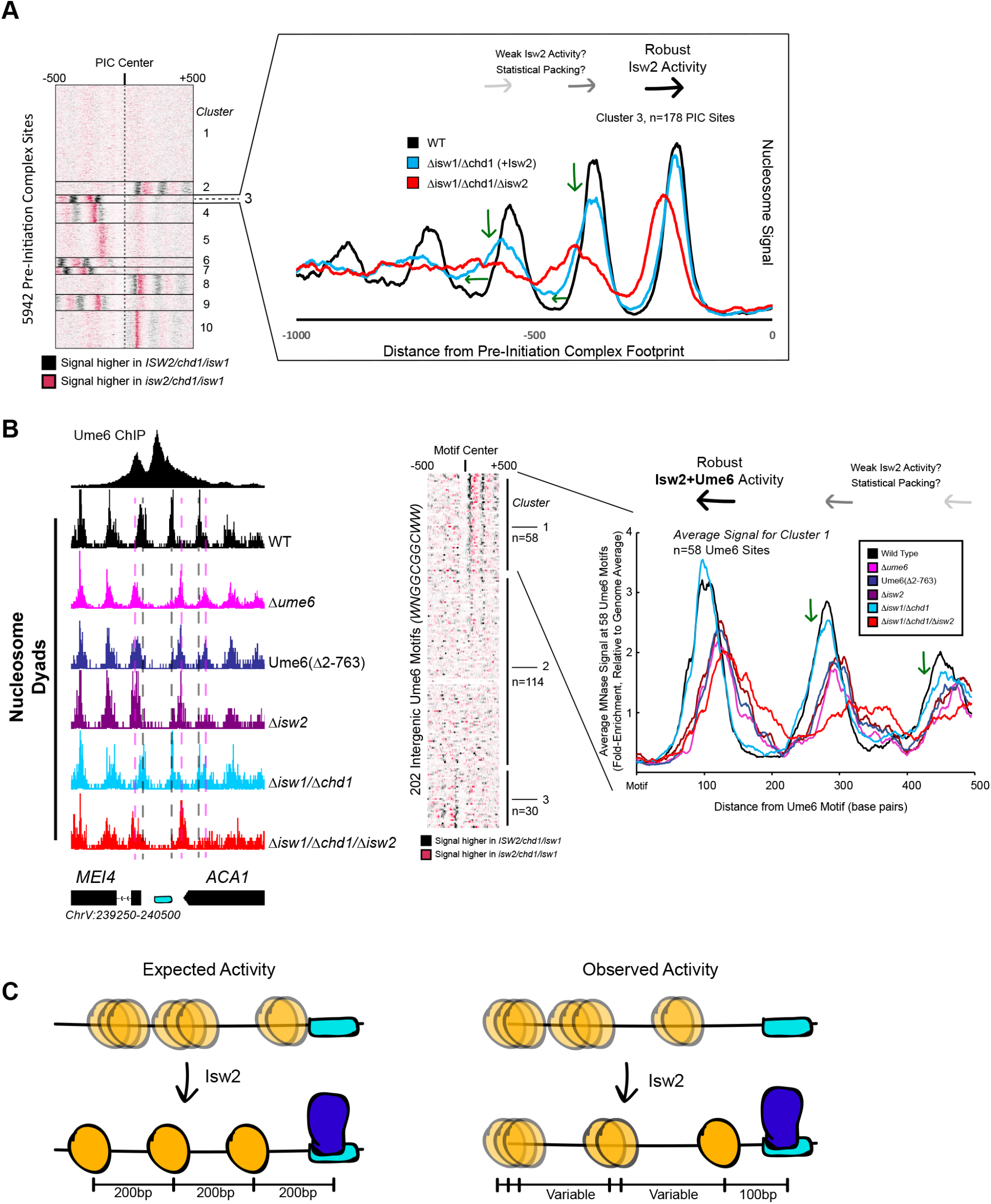
Isw2 is a Specialist Remodeler that Positions Single Nucleosomes at Target Sites. **A** (left) Clustered heat map showing differences in nucleosome dyad signal between *isw2*/*isw1*/*chd1* and *ISW2*/*isw1*/*chd1* strains at 5942 pre-initiation complex sites (PICs). Black indicates positions where Isw2 preferentially positions nucleosomes compared to the strain lacking Isw2. (right) Average nucleosome dyad signal for wild-type (black), *isw1*/*chd1* (cyan) and *isw2*/*isw1*/*chd1* (red) strains for the 178 PIC sites in cluster 3. Black arrows denote Isw2-driven nucleosome shifts. Green arrows indicate rapid decay of positioning at PIC-distal nucleosomes in the *ISW2*/*isw1*/*chd1* mutant. **B** (left) Genome Browser image showing nucleosome dyad signal at a Ume6 motif (cyan rectangle) for indicated strains. Vertical gray dashed line denotes the motif-proximal wild-type nucleosome positions while vertical pink dashed line indicates the nucleosome positions in the absence of Ume6 or Isw2. (center) Clustered heat map showing the difference in nucleosome dyad signal between *isw2*/*isw1*/*chd1* and *ISW2*/*isw1*/*chd1* strains at 202 intergenic Ume6 motifs. Black indicates positions where Isw2 preferentially positions nucleosomes compared to strains lacking Isw2. (right) Average nucleosome dyad signal for indicated strains at Ume6 motifs in cluster 1. Black arrows indicate direction of nucleosome positioning by Isw2. Green arrows signify decreased positioning of motif-distal nucleosomes in the *ISW2*/*isw1*/*chd1* strain (cyan) compared to wild type (black). **C** (left) Cartoon depicting the expected activity of Isw2 at barrier elements according to current biochemical data and nucleosome positioning models. Isw2 is thought to move nucleosomes away from bound factors and space nucleosomes with an approximately 200 base pair repeat length. (right) Cartoon of the observed activity of Isw2 at target sites where only a motif-proximal single nucleosome is precisely positioned but distal nucleosomes are not well-spaced by Isw2.

It has been shown that Isw2 associates with sequence-specific DNA binding factors, such as the transcriptional repressor Ume6^24,25^. Isw2 activity at Ume6-bound loci has been previously characterized as precise, with Isw2 reproducibly moving nucleosomes until the predicted edge of the nucleosome core particle is 30 base pairs from the center of the Ume6 binding motif^29^. Because of the connection to Ume6, we examined nucleosome positions in an *isw1*/*chd1* background in the presence and absence of Isw2 to determine whether Isw2 is similarly restricted at known target sites. Again, we determined that Isw2 is efficient at positioning the Ume6-proximal nucleosome but positioning of nucleosomes decays rapidly as the distance from the proximal nucleosome increases, suggesting that Isw2 may only position single nucleosomes at target sites (Figure 1B). Nucleosomes also appear to always be positioned toward Ume6 motifs, as nucleosome positions in the absence of Isw2 are always more distal to the Ume6 motif than when Isw2 is present. Finally, these nucleosomes are positioned with the dyad only separated from the Ume6 motif by 100 nucleotides, rather than the ~200 nucleotides that would be expected between dyads in a nucleosome array based on Isw2 preferentially leaving 60 base pairs of linker DNA between nucleosomes *in vitro^9,11^*.

The observations that 1) Isw2 is solely required to move single nucleosomes at target sites, 2) Isw2 does not have global nucleosome spacing/organizing activity and 3) Isw2 moves nucleosomes within 100 nucleotides of bound Ume6 suggest that Isw2 behavior in cells is distinct from our understanding of Isw2 activity from decades of biochemical characterization. Similarly, these specific movements toward Ume6 (a barrier) are inconsistent with previous biophysical studies, where ISWI proteins were shown to move nucleosomes away from inert DNA-bound factors^37^. Because of these inconsistencies, we wished to know if Isw2 followed the “barrier model” for positioning nucleosomes at Ume6-bound targets. To initially test this, we created a variant Ume6 construct where all residues were deleted except for the DNA binding domain. This Ume6(Δ2-763) construct binds to the same targets as full-length Ume6 (Figure S1D). However, Isw2 does not appear to have any activity on global Ume6-proximal nucleosomes in the presence of the Ume6 DNA binding domain alone, as nucleosomes in this strain occupy identical positions as when Ume6 or Isw2 are completely absent (Figure 1B). In the presence of full-length Ume6, the Isw2 complex appears to be necessary and sufficient for moving motif-proximal nucleosomes, as nucleosome positions in the *ISW2/isw1/chd1* strain could achieve identical motif-proximal nucleosome positions as the wild-type strain. Additionally, the *CHD1/isw1/isw2* and *ISW1/chd1/isw2* strains were unable to move any Ume6-proximal nucleosomes (Figure S1E,F), which strongly argues that Ume6 is not acting as a passive barrier against which nucleosome spacing factors can pack nucleosomes. Instead, these data are more consistent with the recent characterization of Isw2 as a “puller”^38^, with Ume6 being a DNA-bound factor that may immobilize Isw2 to create leverage for “pulling”. Consistent with this immobilized pulling model and consistent with the directional movement of single nucleosomes toward Ume6-bound sites, artificially tethered chromatin remodeling proteins were previously shown to always move nucleosomes toward target sites^27^. We suspected that Ume6 and Isw2 likely interact in a specific fashion to faithfully select and precisely move single target nucleosomes toward a recruitment motif (Figure 1C).

### A Small Helical Epitope is Necessary and Sufficient for Isw2-Directed Nucleosome Positioning at Ume6 Targets

To determine which region(s) on Ume6 are required for specific nucleosome positioning by Isw2, we initially created a panel of N-terminal Ume6 truncations to determine when nucleosome positioning by Isw2 is lost (Figure S2A). This initial truncation panel was necessary due to the poor overall conservation of the Ume6 protein even within related yeasts, as well as the disordered structure predicted by Phyre2^39^. Our truncation panel indicated that Isw2 activity was retained if the N-terminus was deleted to residue 322 but lost when deleted to residue 508. Closer inspection of the residues between 322 and 508 revealed a conserved region with a proline-rich segment followed by a predicted alpha helix, altogether spanning Ume6 residues 479-508 (Figure 2A). Deletion of residues 2-479 preserved Isw2-positioned nucleosomes at Ume6 sites, while an internal deletion of 480-507 in the context of an otherwise full-length Ume6 abrogated nucleosome positioning by Isw2 (Figure 2A, S2B). Importantly, Ume6 Δ2-479 and Ume6 Δ2-508 showed identical binding as measured by ChIP (Figure S2C) indicating that the loss of nucleosome positioning is not due to loss of Ume6 binding.

Since this region is proximal to the characterized Sin3-binding domain in Ume6^40^, we wished to validate that the newly determined Isw2 recruitment helix is independent from the Sin3-binding domain. Ume6 recruits both Isw2 and Sin3-Rpd3 for full repression of target genes^24,25^. If either Isw2 or Sin3-Rpd3 is present, there is partial repression at Ume6-regulated genes. However, if Sin3-Rpd3 and Isw2 are both lost, Ume6 targets are fully de-repressed. We examined transcriptional output at Ume6 genes in Ume6(Δ2-479) +/− Rpd3 and Ume6(Δ2-508) +/− Rpd3. Transcription was modestly increased at Ume6 targets in Ume6(Δ2-508)/*RPD3+* compared to Ume6(Δ2-479)/*RPD3+* (Figure S3A), which would be expected if only Isw2 is lost when residues 479-508 are deleted. More convincingly, only a modest increase in transcription was seen at Ume6 targets in the Ume6(Δ2-479)/Δ*rpd3* strain suggesting Isw2 is still present, while the Ume6(Δ2-508)/Δ*rpd3* strain displayed extreme induction of Ume6-regulated genes, suggesting that both Isw2 and Rpd3 activity are absent (Figure 2B, S3B).

Finally, we wanted to know if the predicted helix consisting of Ume6 residues 479-508 was sufficient to bring Isw2 nucleosome positioning activity to Ume6 target sites. To test this, we employed the SpyCatcher/SpyTag system^41^, which creates a spontaneous covalent bond between a short SpyTag peptide and a SpyCatcher domain. We fused the SpyTag peptide to the C-terminus of Ume6(Δ2-596), a construct that is incapable of positioning motif-proximal nucleosomes (Figure S2A). We then appended Ume6 residues 479-508 to the C-terminus of the SpyCatcher domain and introduced this fusion on a yeast expression plasmid driven by the ADH1 promoter. In yeast cells, this would create a fusion protein where the helical element is ectopically displayed on the C-terminus of a DNA-binding competent but nucleosome-positioning deficient construct, connected via a SpyTag-SpyCatcher linker. This fusion protein was capable of fully recapitulating Isw2-positioned nucleosomes at a subset of Ume6 sites (Figure 2C, S4A, S4B). Perhaps not surprisingly, considering the non-native positioning of the recruitment helix in this fusion construct, not all Ume6 sites were able to gain proper nucleosome positioning with this chimeric system (Figure S4A). We conclude that the region spanning residues 479-508 in Ume6 is a yeast-conserved Isw2-recruitment domain and is required and sufficient for recruiting Isw2 nucleosome positioning activity to Ume6 targets.

### A Similar Helical Element Exists in Swi6, a Newly Identified Isw2-Recruitment Adapter Protein

While dissecting the Isw2 recruitment domain in Ume6, we discovered that deleting the *MBP1* gene resulted in ectopic nucleosome positioning at a subset of Mbp1 target loci, which was identical to mispositioned nucleosomes in a Δ*isw2* strain. Mbp1 is a conserved cell cycle regulator that complexes with Swi6 to form the MBF complex^42^. This complex activates the transition from G1 to S and includes the conserved function of regulating Start-specific transcription^42,43^. To determine how Mbp1 recruits Isw2, we similarly made truncations of Mbp1 to determine at which point nucleosome positioning no longer resembles wild-type positioning and reflects Δ*isw2* positioning instead. The DNA binding element in Mbp1 resides in the extreme N-terminus (Figure 3A) spanning residues 2-124^44^, so a panel of C-terminal truncations was created. However, before examining the full panel of truncations, we observed that nucleosome positioning was already identical to Δ*isw2* positioning in Mbp1 Δ562-833, the first C-terminal truncation examined (Figure S5). This extreme C-terminal region interacts with Swi6 (Figure 3A), so we speculated that Swi6 may be responsible for recruiting Isw2. As predicted, deletion of the *SWI6* gene led to ectopic nucleosome positions identical to Δ*mbp1* and Δ*isw2* strains at the small subset of Mbp1 targets.

**Figure 2.**
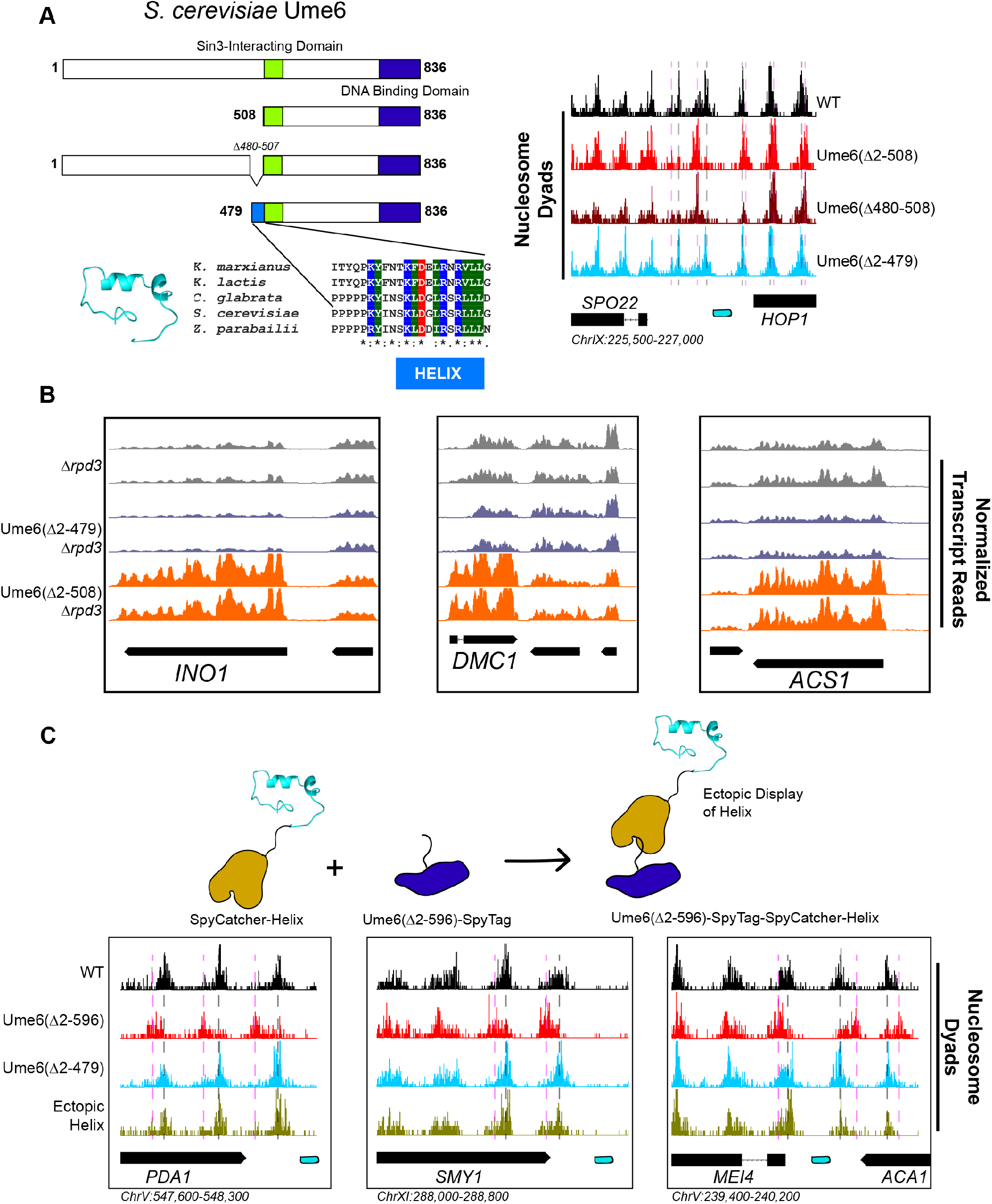
A Small Predicted Helix is the Isw2-Recruitment Epitope in Ume6. **A** (top left) Schematic diagram of Ume6 truncation and deletion constructs used to identify the Isw2 recruitment epitope, with the known Sin3-Interacting Domain depicted as a green square, the DNA binding domain as a dark blue rectangle, and the putative Isw2-recruitment helix as a light blue rectangle. (bottom left) Modeled helical peptide (by Phyre2) and sequence conservation of the identified Isw2-recruitment motif in Ume6 constructs from other yeasts. Asterisks denote invariant residues. (right) Nucleosome dyad signal for Ume6 truncation and deletion strains indicate deletion of the region from residue 480-507 completely abrogates nucleosome positioning by Isw2 at Ume6 target sites. Vertical dashed gray lines denote wild-type positions of nucleosomes, while vertical dashed pink lines indicate *isw2* or *ume6*-deficient positions of nucleosomes. **B** Genome Browser image showing transcript abundance at three Ume6 target sites for yeast strains lacking Rpd3 with wild-type Ume6 (gray), Ume6(Δ2-479) (blue) and Ume6(Δ2-508) (orange). Grossly increased transcription is seen when residues 480-507 are deleted, consistent with expected transcriptional increase associated with loss of Isw2 and Rpd3. No significant increase in transcription is detected when Ume6 residues 2-479 are deleted. Biological replicates are shown to highlight reproducibility. **C** (top) Cartoon schematic for ectopic display of the Isw2-recruiting helix (residues 480-507) to the C-terminus of a truncated Ume6 construct lacking Isw2-directed nucleosome positioning. A short SpyTag is appended to the C-terminus of the Ume6 construct and residues 480-507 are fused to the SpyCatcher domain and introduced on a yeast expression vector. (bottom) Nucleosome dyad signal demonstrating recovery of Isw2-directed nucleosome positions at a subset of Ume6 target genes by the ectopically displayed helical element. Vertical dashed gray lines denote wild-type positions of nucleosomes, while vertical dashed pink lines indicate *isw2* or *ume6*-deficient positions of nucleosomes. Individual biological replicates for nucleosome positions after ectopic display of the recruitment helix are provided in Figure S4.

We conducted sequence alignment and conservation analyses between the helical element in Ume6 and full length Swi6 from multiple yeast species (Figure 3A). We noticed a similarly conserved surface-exposed helix^45^ in the cell cycle regulating protein Swi6 (Figure 3A). Intriguingly, the function of this helical element has not been determined despite its sequence conservation. Because Swi6 also interacts with Swi4 to form the highly-conserved SBF complex^42^, we speculated that deletion of either Swi6, Swi4 or Isw2 could potentially lead to ectopic nucleosome positions at a subset of SBF targets. Indeed, we observed ectopic nucleosome positioning at the HSP12 locus (an SBF target) when either Isw2, Swi6 or Swi4 was absent (Figure 3A). Wild-type nucleosome positions were observed in the absence of Mbp1, indicating that this is specific to SBF. Similarly, wild-type nucleosome positions were observed at Mbp1 targets when Swi4 was missing (Figure 3A), again suggesting that MBF and SBF have individual Isw2-targeting capacity at their respective binding sites. Swi6 appears to be an adapter protein responsible for recruiting Isw2 to Mbp1 and Swi4 sites, since Swi6 has no intrinsic DNA binding domain.

**Figure 3.**
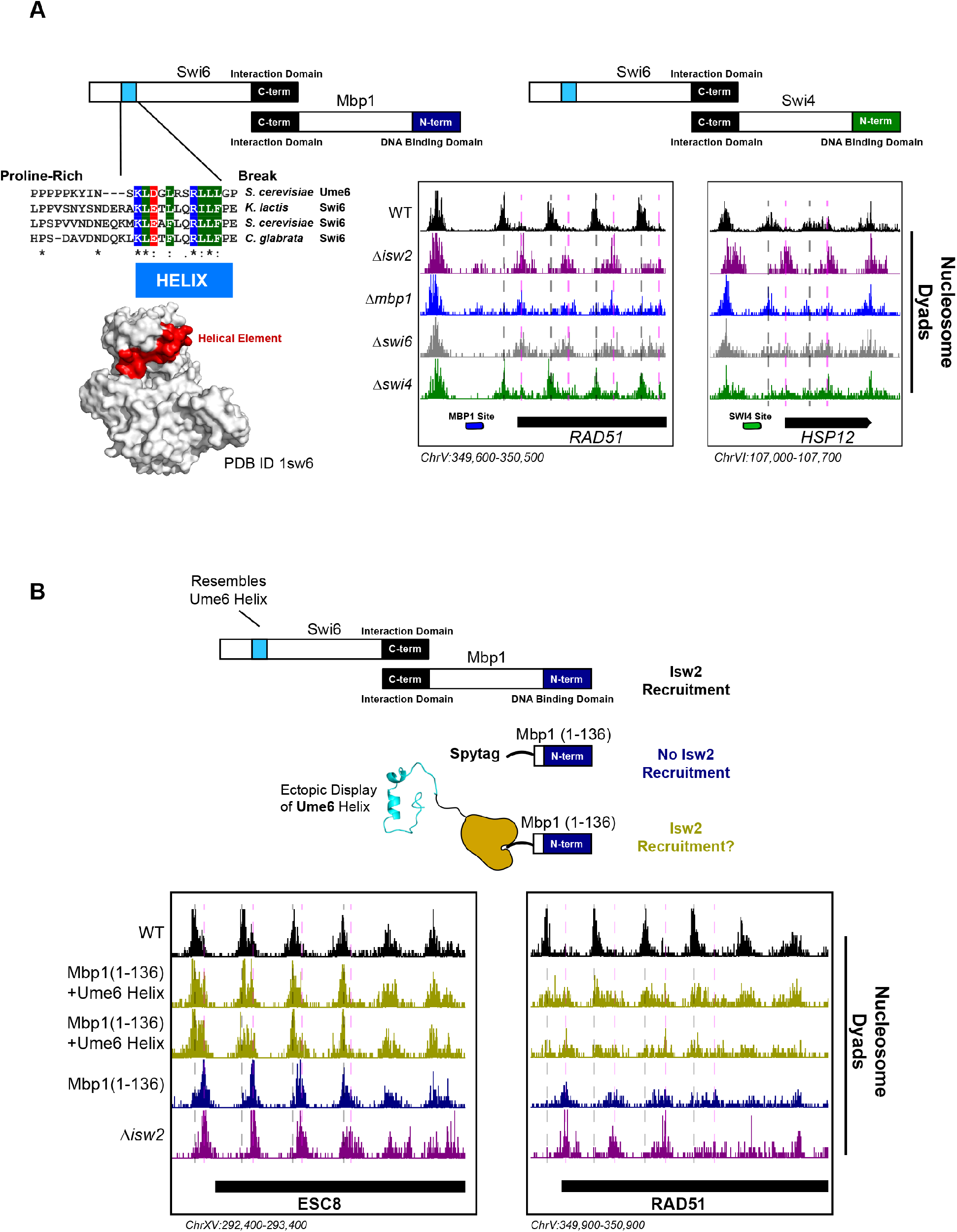
The Cell Cycle Regulator Swi6 Contains a Similar Helical Element and Recruits Isw2 to MBF and SBF target genes. **A** (top left) Schematic representation of the Swi6-Mbp1 MBF complex. Swi6 interacts with Mbp1 through the C-terminal domain (black rectangle). Mbp1 has an N-terminal DNA binding domain (dark blue rectangle). The putative Isw2 recruitment helix is in the Swi6 N-terminus (light blue rectangle). (center left) Conserved residues in the putative Isw2-recruitment helix in Swi6 for three yeast species compared to the Isw2-recruitment helix in Ume6 for *S. cerevisiae*. (bottom left) Crystal structure (PDB ID 1sw6) showing the location of the surface-exposed, conserved helical element from Swi6 in red. (top right) Schematic representation of the Swi6-Swi4 SBF complex. Swi6 interacts with Swi4 through the C-terminal domain (black rectangle). Swi4 has an N-terminal DNA binding domain (green rectangle). Putative Isw2-recruitment helix is shown (small blue rectangle) (bottom center) Genome Browser image showing nucleosome dyad signal for indicated strains at the *RAD51* locus, an MBF target gene with an indicated Mbp1 binding motif (blue rectangle). Wild-type nucleosome positions are indicated by vertical dashed gray lines while ectopic positions associated with *isw2*, *mbp1*, and *swi6* deletion strains are indicated by vertical dashed pink lines. (bottom right) Genome Browser image showing nucleosome dyad signal for indicated strains at the *HSP12* locus, an SBF target gene with an indicated Swi4 binding motif (green rectangle). Wild-type positions are denoted by vertical gray dashed lines while ectopic nucleosome positions associated with *isw2*, *swi6* and *swi4* deletion strains are indicated with vertical pink dashed lines. **B** (top) Schematic representation of constructs used to determine if ectopic display of an Isw2 recruitment helix on the Mbp1 N-terminus could recover Isw2-positioned nucleosomes at Mbp1 target genes. Either wild-type Mbp1, a C-terminal deletion of Mbp1 leaving only the DNA binding domain and an appended SpyTag, or a C-terminal deletion of Mbp1 leaving the DNA binding domain and SpyTag with constitutively-expressed SpyCatcher fused to the Isw2-recruitment helix from Ume6 were examined. (bottom) Genome Browser image showing nucleosome dyad signal for indicated strains at the *ESC8* (left) or *RAD51* (right) loci. Gray vertical dashed lines indicate wild-type nucleosome positions while vertical dashed pink lines indicate ectopic nucleosome positions associated with inactive Isw2 or Mbp1/Swi6. Biological replicates for ectopic display of the recruitment helix are provided as two separate tracks (gold) to emphasize reproducibility.

To determine if Isw2 recruitment to Mbp1 sites was sufficient to recapitulate proper nucleosome positioning, we again used a SpyTag-SpyCatcher approach (Figure 3B). Mbp1 was truncated to the DNA binding domain alone (Mbp1 1-136), which abolishes its interaction with Swi6 but still allows for proper genomic localization. This truncation construct was appended with SpyTag, and nucleosome positions were examined in the absence of any SpyCatcher partner present. As expected, we observed aberrant chromatin structure identical to the Δ*isw2* strain near the Isw2-dependent Mbp1 targets, adjacent to Mbp1 consensus motifs (Figure 3B). We then introduced SpyCatcher fused to the helical element from Ume6, which was characterized above for bringing Isw2 to Ume6-bound loci. Introduction of the SpyCatcher-Ume6 fusion to the Mbp1(1-136)-SpyTag background resulted in the rescue of proper Isw2-directed nucleosome positioning at Mbp1 sites (Figure 3B). Altogether, these data strongly support our model that these conserved, putatively helical sequences are important for recruiting Isw2 to establish proper chromatin structure at multiple sequence-specific motifs throughout the genome. We also implicate Swi6 as an adapter protein for bringing Isw2 to a small subset of both Swi4 and Mbp1 targets to create Isw2-specific nucleosome positioning at these genes. Finally, the ectopic display of an Isw2-recruitment helix can recapitulate proper Isw2-directed nucleosome positioning, further supporting the notion that a small epitope is necessary and sufficient for communicating specific nucleosome positioning outputs to the Isw2 chromatin remodeling protein.

### The Conserved WAC Domain in Itc1 is the Targeting Domain of the Isw2 Complex

The Isw2 complex contains two major subunits (Figure 4A). The catalytic subunit Isw2 harbors the energy-producing ATPase domain flanked by biochemically well-defined autoregulatory domains^46–48^ with a C-terminal HAND-SANT-SLIDE (HSS) domain, thought to bind linker DNA^34^ and interact with the accessory subunit Itc1. Itc1 contains an N-terminal WAC domain, thought to bind to and sense extranucleosomal DNA and help with nucleosome assembly in the Drosophila ortholog ACF1^49^. Itc1 links to Isw2 through a DDT domain^49^. The ~350 amino acid N-terminal region of human Acf1 was shown to bind both extranucleosomal linker DNA and the histone H4 tail, suggesting an allosteric mechanism through which ISWI complexes can set proper spacing between nucleosomes^50^. Though this work was performed with human ACF complex, Hwang *et al* demonstrated removal of residues 2-374 in *S. cerevisiae* was lethal, suggesting a critical and conserved role of these residues in establishing proper chromatin structure *in vivo*^50^.

**Figure 4.**
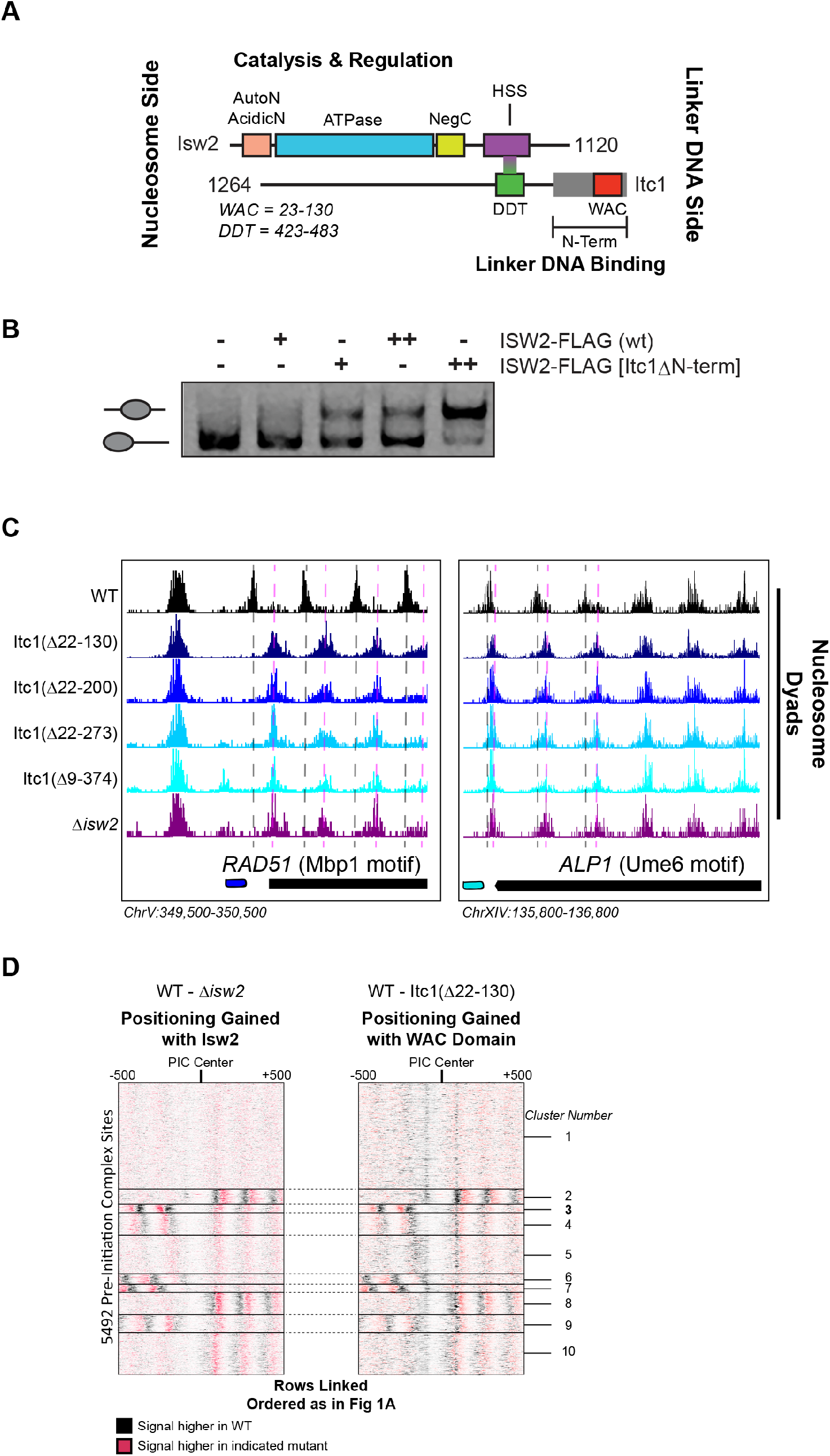
The N-terminal WAC Domain in Itc1 Couples Isw2 Biochemical Activity to all Isw2 Genomic Targets. **A** Cartoon representation the Isw2 and Itc1 subunits of the yeast ISW2 complex. Isw2 possesses autoregulatory domains on either side of the catalytic ATPase domain (AutoN and NegC). The HAND-SANT-SLIDE domain of Isw2 interacts with the DDT domain of Itc1 for complex formation. Itc1 has an N-terminal region thought to act as a length-sensing domain (gray rectangle) and an N-terminal WAC domain with putative nonspecific linker DNA binding ability. **B** Nucleosome sliding assay demonstrating that deletion of the N-terminal domain (Δ9-374) from Itc1 does not impair nucleosome sliding *in vitro* by the Isw2 complex. Higher electrophoretic mobility indicates end-positioned (unslid) nucleosomes while lower electrophoretic mobility indicates centrally positioned (slid) nucleosomes. Isw2-FLAG complexes were purified from exponentially growing yeast cells as described in the methods. Amount of Isw2 added was 1 μl (+) or 1.5μl (++). Sliding assays were performed 3 independent times with similar results. **C** Genome Browser images showing nucleosome dyad positions for indicated strains at *RAD51* and *ALP1*, two representative Isw2 targets. Only wild-type cells display the proper nucleosome positions (vertical gray dashed lines) while all Itc1 truncations and *isw2* deletion display similar ectopic nucleosome positions (vertical pink dashed lines). **D** Heat map comparing difference in nucleosome positions at 5942 PIC locations for *isw2* deletion versus wild-type strains (left) and Itc1(Δ22-130) versus wild-type strains (right). Black indicates where nucleosomes are shifted by functional Isw2 while red indicates where nucleosomes shift when Isw2 complex is perturbed. All rows are linked and ordered identically to Figure 1A.

Because of the geometry of the Isw2 complex, with the N-terminus of Itc1 sensing DNA information distal to the nucleosome onto which the catalytic subunit is engaged, we speculated that the N-terminus of Itc1 would be the most likely component of the Isw2 complex for interacting with epitopes in DNA-bound recruitment factors. We first attempted to recapitulate the result from Hwang *et al* and made the identical Itc1(Δ2-374) deletion. Isw2 containing Itc1(Δ2-374) did not display any defects in nucleosome sliding using a gel mobility shift assay that detects nucleosome centering by Isw2 (Figure 4B). Surprisingly, this construct was not lethal in our W303 background, but phenocopied a Δ*isw2* strain by displaying identical ectopic nucleosome positioning at all Isw2 target sites throughout the genome (Figure 4C). Since proper targeted nucleosome positioning was lost when this large N-terminal region was removed, but complex formation and catalytic activity were maintained, we strongly suspected that the Isw2 targeting domain resided in the Itc1 N-terminus. We created a panel of truncations in this region, guided by sequence conservation through humans, and determined whether wildtype or Δ*isw2* positions were observed throughout the genome. All truncations tested resulted in loss of positioning at Isw2 targets, and we were able to narrow the targeting region entirely to the highly conserved WAC domain. Deletion of the WAC domain (Itc1 residues 22-130) produced identically ectopic nucleosome positions compared to Δ*isw2* at target loci (Figure 4C) and genome-wide (Figure 4D). We conclude that the WAC domain of Itc1 is the component of the Isw2 complex responsible for coupling with epitopes on DNA-bound factors such as Ume6, Swi6, and all other Isw2 targeting proteins with yet-to-be-defined recruitment epitopes.

### The WAC Domain Binds Isw2 Targets and Orients the Catalytic Subunit on Target-Proximal Nucleosomes

To confirm that the WAC domain can interact with Isw2 targets throughout the genome, we created Itc1(1-73)-FLAG and Itc1(1-130)-FLAG constructs based on two differentially conserved regions within the full WAC domain (Figure 5A). Neither of these constructs contains the DDT domain, so they are incapable of forming a complex with endogenous Isw2. We performed ChIP-Seq to determine if these WAC domain constructs could associate with Isw2 targets without complexing with the Isw2 catalytic domain (Figure 5B, S6). Genome-wide binding demonstrates large, but not complete overlap of Isw2(K215R)-FLAG ChIP peaks with both Itc1(1-73)-FLAG and Itc1(1-130)-FLAG, strongly suggesting that the Itc1 region from 1-73 alone can interact with Isw2 targets.

**Figure 5.**
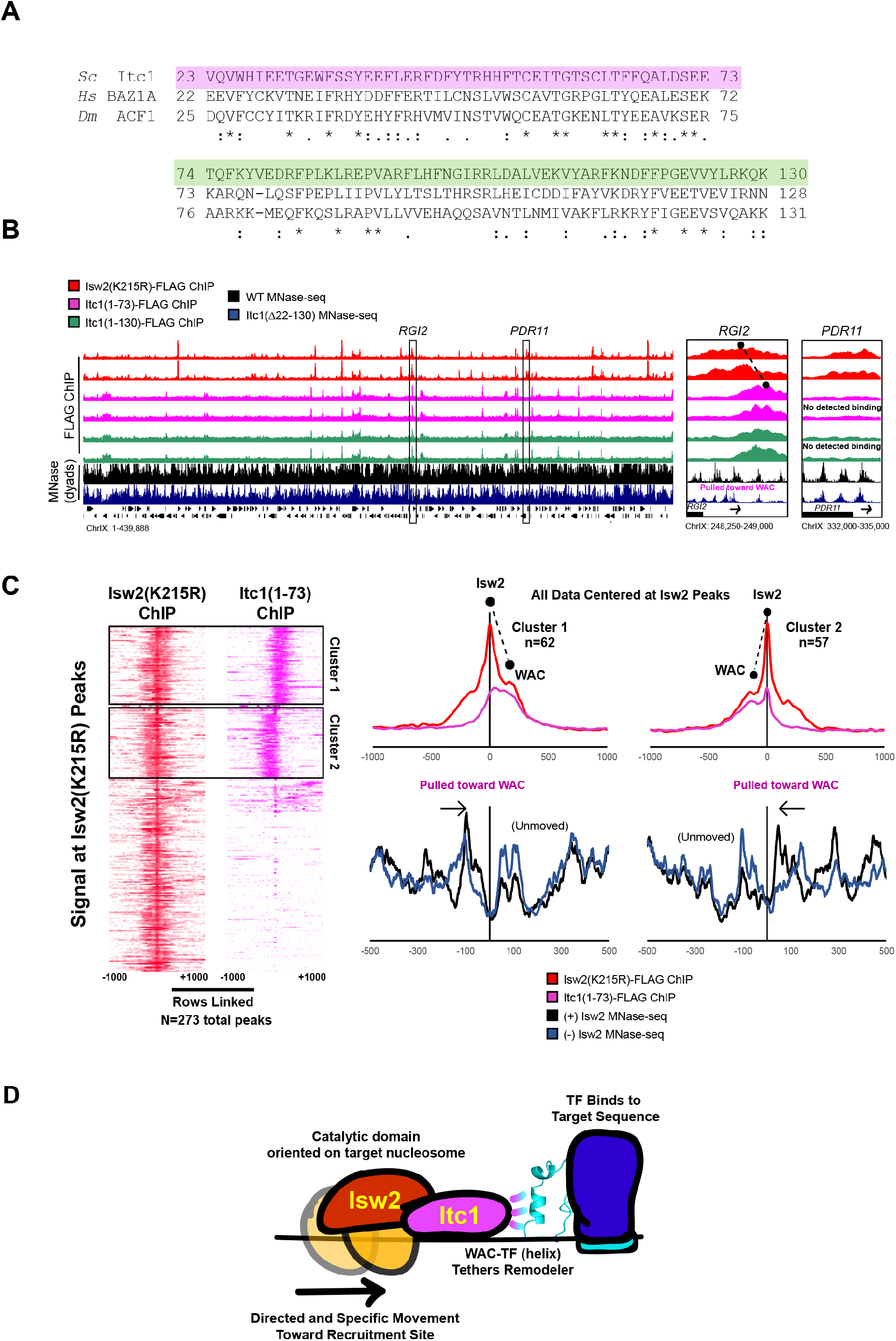
The Itc1 WAC Domain Associates with Genomic Isw2 Targets and Orients Isw2 on the Proper Nucleosomes. **A** Sequence conservation for regions of Itc1 examined by ChIP. Itc1(1-73)-FLAG incorporates the pink highlighted region while Itc1(1-130)-FLAG incorporates the pink and green highlighted regions. Sequence conservation is shown relative to human BAZ1A and *Drosophila melanogaster* Acf1, two widely studied Itc1 orthologs. **B** (left) Full view of yeast chromosome IX showing Isw2(K215R)-FLAG ChIP (red), Itc1(1-73)-FLAG ChIP (pink), Itc1(1-130)-FLAG ChIP (green), nucleosome dyad signal from wild-type yeast (black) and nucleosome dyad signal from Itc1(Δ22-130) yeast (blue). Regions indicated by black rectangles are shown with higher resolution on the right. (right) Zoomed-in view of a locus where Isw2-ChIP and Itc1 truncation ChIP overlap (*RGI2*) or where only Isw2 binding is detected (*PDR11*). Black circles indicate center of ChIP peaks and are connected by a dashed black line to highlight offset of indicated peaks. **C** (left) Heat map showing 273 detected Isw2 ChIP peaks (red) clustered by associated Itc1(1-73)-FLAG ChIP (pink). The two clusters (right-side Itc1 and left-side Itc1) are shown on the right. (right) Meta-analysis of Isw2(K215R)-FLAG ChIP signal at 62 cluster 1 peaks or 57 cluster 2 peaks (from left) with associated Itc1(1-73)-FLAG signal. The offset between Isw2 and Itc1 is indicated by two circles connected by a dashed line. Associated nucleosome positions for wild-type and *isw2* deletion strains for each cluster are shown below in black and blue, respectively. All data are centered at called Isw2 peaks. **D** Cartoon representation for how the N-terminal WAC domain of Itc1 interacts with a helical element in a sequence-specific DNA-associated transcription factor to orient Isw2 on the proper motif-proximal nucleosome for directional movement toward the recruitment site.

We noticed that the Itc1 signal and Isw2 signal were offset at target genes such that Itc1(1-73) or Itc1(1-130) was upstream and Isw2 was closer to the nucleosome that was selected for repositioning (Figure 5B, S6). Genome-wide analysis showed that Itc1(1-73) was associated with approximately half of Isw2-bound loci and was offset from the catalytic subunit at all co-bound sites (Figure 5C). In all cases, Itc1(1-73) was found upstream of the nucleosome that was repositioned, and Isw2 was located on top of the selected nucleosome. Nucleosomes were always shifted toward the Itc1 subunit (Figure 5C). This geometry matches what was seen by ChIP-Exo mapping with Isw2 subunits at Reb1 target sites^51^. We propose a mechanism where the Itc1 WAC domain interacts with a DNA-bound factor, which constrains the Isw2 catalytic subunit to select the proper proximal nucleosome and reposition it toward the immobilized Itc1 (Figure 5D). This is again consistent with the recently proposed “pulling” model^38^, but we postulate that Itc1 is anchored to a DNA-bound factor such as Ume6 to allow Isw2 to pull nucleosomes toward the proper location.

### Essential Acidic Residues Required for Targeting are Lost in the Drosophila Genus, Explaining Biochemical and Genetic Inconsistencies

There is an abundance of literature suggesting that *Drosophila* ACF complex, the Isw2 ortholog, is a nonspecific nucleosome spacing and assembly factor that evenly spaces phased nucleosome arrays against defined genomic barriers^10,18,49^. We wondered if the WAC domain of *Drosophila* Acf1 was different from that of Itc1, so we performed sequence alignment of WAC domains and compared to Acf1 from the *Drosophila* genus. While sequence alignment demonstrated widespread conservation of the WAC domain, one striking feature was exposed: the *Drosophila* genus underwent reversal or loss of negative charge at multiple residues that are strictly or mostly acidic in other representative organisms (Figure 6A).

**Figure 6.**
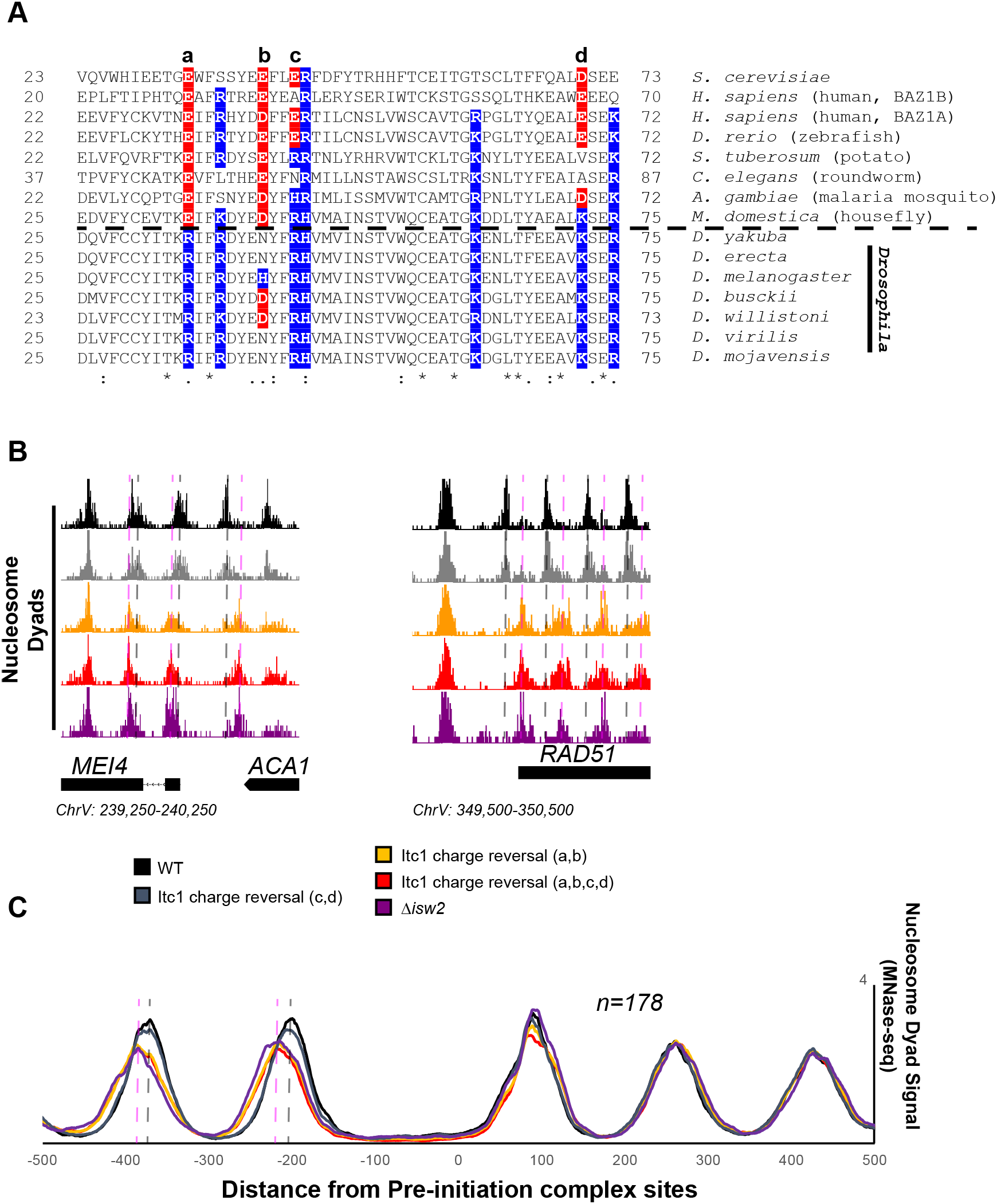
Essential Targeting-Specific Charged Residues in the Conserved WAC Domain is Lost in *Drosophila*. **A** Sequence conservation of the N-terminal region (22-73) of Itc1 across various organisms with key charged residues highlighted in blue or red for positive and negative charge, respectively. Horizontal dashed line indicates separation of all other species from members of the *Drosophila* genus. **B** Genome Browser image showing nucleosome dyad signal at two representative Isw2 target loci. Wild-type nucleosome positions are indicated by gray vertical dashed lines while ectopic nucleosome positions associated with *isw2* deletion or indicated charge reversal mutations are denoted by vertical pink dashed lines. **C** Meta-analysis of nucleosome dyad signal at 178 PIC sites associated with cluster 3 (from Figure 1A). Only wild type and charge reversal c,d display proper nucleosome positions while charge reversal a,b or a,b,c,d display ectopic positions identical to deletion of *ISW2* completely.

Two of these residues are strictly acidic in all organisms except members of the *Drosophila* genus (E33 and E40 in Itc1). The other two (E43 and D70 in Itc1) are more loosely conserved, though they are strictly positive charge in *Drosophila*. We made charge-reversal mutations in *S. cerevisiae* Itc1 to recapitulate the *D. melanogaster* residues at each of these positions either pairwise (a,b and c,d to separate the strictly-conserved acidic versus loosely conserved acidic nature) or simultaneously (a,b,c,d) to reverse all charges to the *D. melanogaster* sequence. We assessed whether charge reversal was sufficient to abrogate targeted nucleosome positioning at Isw2 targets across the yeast genome (Figure 6B). Strikingly, the E33R/E40H double mutation (a,b) was enough to completely abolish Isw2 activity at specific and known Isw2 targets (Figure 6B) and at all genomic loci where Isw2 activity is observed (Figure 6C). Mutation of the less-conserved acidic residues E43R/D70K (c,d) retained Isw2-directed nucleosome positioning. As expected, mutation of all four acidic residues (a,b,c,d) E33R/E40H/E43R/D70K resulted in complete loss of Isw2 targeted activity across the genome (Figure 6B,C). We conclude that the *Drosophila* genus lost critical acidic residues that are essential for targeted nucleosome positioning by *S. cerevisiae* Isw2, potentially explaining the disconnect between the *Drosophila* ACF literature and what we have characterized herein. It is possible that the increase in positive charge simultaneously increases nonspecific binding of *Drosophila* Acf1 to extranucleosomal DNA, and these charge reversals may help explain the nonspecific spacing behavior of Acf1 observed in *Drosophila*. We also believe there is strong potential that humans and most other organisms have retained targeting potential, as they retain mechanistically important acidic residues present in yeast Itc1. In support of conservation, targeted nucleosome array formation has previously been observed in humans at specific transcription factor sites including CTCF, Jun and RFX5^17^.

## Discussion

### An Interacting Barrier Model for Nucleosome Array Establishment

Collectively, our results give rise to an “interacting barrier model” as an alternative means of genomic nucleosome positioning by introducing a targeted interaction between an epitope contained within conditionspecific transcription factors and ISWI-type ChRPs (Figure 7). We show that a recruitment factor, the sequence-specific repressor Ume6, harbors a helical domain that interacts with the N-terminus of the Isw2 accessory protein Itc1. Further, we reveal this geometrically restricts the binding of the Isw2 catalytic subunit to a motif-proximal nucleosome. The complex then remodels the nucleosome, repositioning it to a specific distance from the Ume6 recognition motif. At this point and for reasons to be elucidated, this complex is strained or inactivated, and it fails to remodel any further, leaving the nucleosome in a precise location with respect to the bound recruitment factor. The activity of Isw2 and the interacting barrier sets the absolute phase of a nucleosome array that is propagated by true nonspecific spacing activities of Chd1 and Isw1 in yeast, as previously described^19,20,36^. This “interacting barrier model” of chromatin organization is more comparable to the factor-targeted activities of SWI/SNF than the non-specific array spacing of CHD family remodelers, and is potentially conserved through humans based on conservation of key interacting residues in Itc1 (Figure 6A) and the observation that Isw2 orthologs can precisely position nucleosomes adjacent to specific factors in the human genome^17^. Together, we show coupling between an epitope on an interacting barrier and a conserved chromatin remodeling protein leads to robust, directional and specific nucleosome organization at genomic regulatory elements.

**Figure 7.**
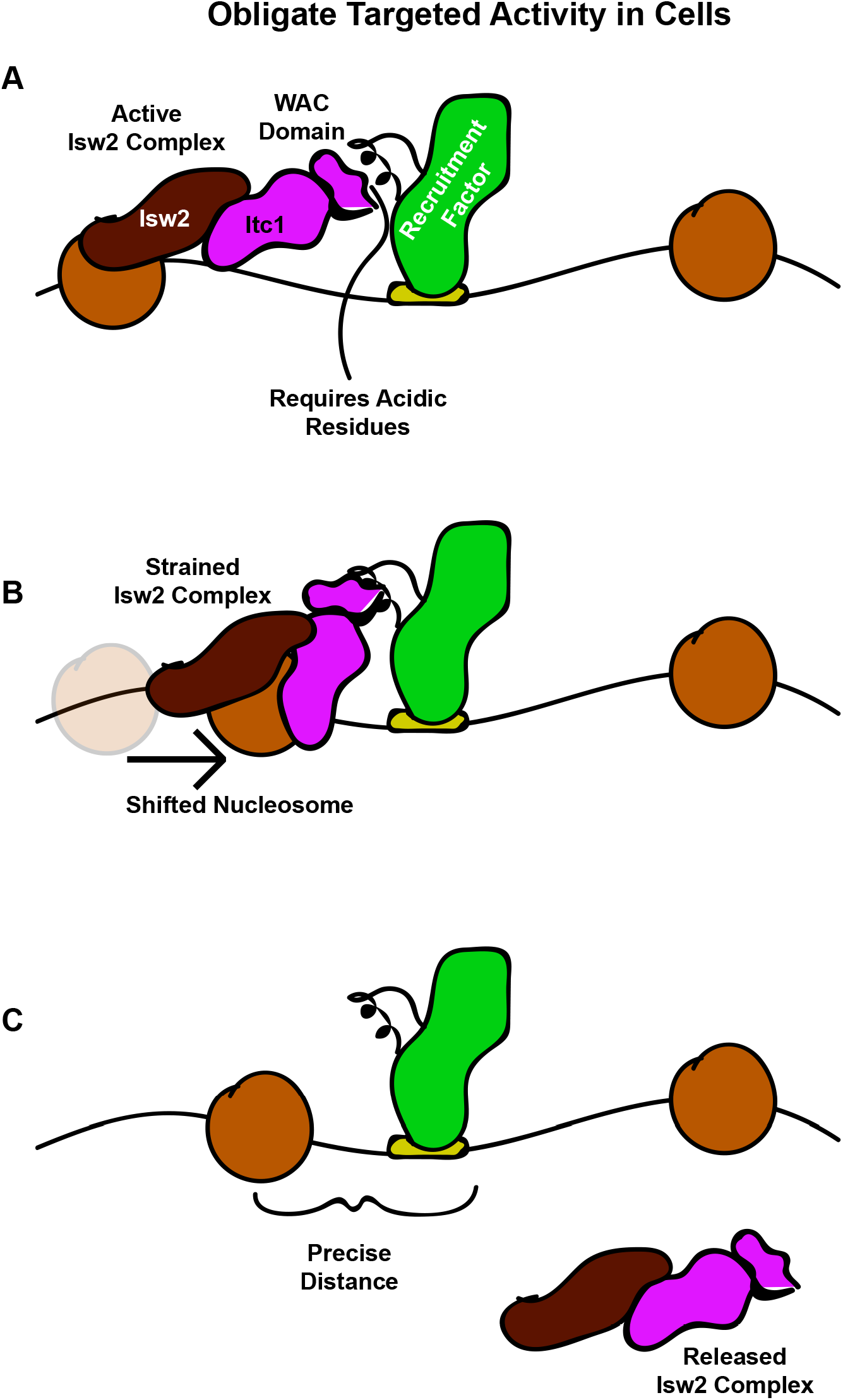
The Interacting Barrier Model for Specific Nucleosome Placement by Isw2. **A** A DNA binding factor with an Isw2-recruitment helix (or other epitope) associates with DNA. The WAC domain of Itc1 engages with the recruitment epitope to proximally-align the catalytic subunit of Isw2 with the proper nucleosome. **B** Nucleosome sliding by Isw2 creates a nucleosome position that is too close to the recruitment epitope for proper alignment of the recruitment epitope - Itc1 WAC - Isw2 catalytic subunit axis, leading to a “strained complex”. **C** The precise distance between the DNA-bound Isw2 recruitment factor and proximal nucleosome after nucleosome positioning by Isw2 is no longer a good substrate for Itc1 WAC interaction and further remodeling, so the Isw2 complex diffuses to new target loci.

### Small Epitopes in Transcription Factors Organize Large Chromatin Domains

Our data suggest that some small peptide domains embedded within transcription factors can nucleate nucleosome arrays of over 1kb in length *in vivo* through an interaction with evolutionarily conserved ChRPs. Unlike the arrays established by non-specific ChRPs, these nucleosome arrays are organized in a sequence-specific and directional manner. Establishing large swaths of chromatin structure by appending a small epitope on a genome-associated protein creates opportunity for diversity with few evolutionary constraints. Only changes in relatively small DNA binding motifs and the small peptide sequences with which they interact can have large impact on chromatin structure. Supporting this notion, we were able to identify a strikingly similar motif to that found in Ume6 in the unrelated cell cycle regulator Swi6, which we identified as a new Isw2 recruitment adapter for Swi4 and Mbp1. For these reasons, we find it likely that more ChRP-interacting motifs will be discovered in multiple transcription factors from a variety of organisms, and these motifs may play a significant role in sequence-specific nucleosome positioning for precisely phased and tunable nucleosome arrays in eukaryotic genomes. Importantly, the identification of such epitopes in human cells could lead to the development of targeted drugs to specifically disrupt defined remodeler-transcription regulator interactions.

### Isw2 is Obligately Targeted to Specific Nucleosomes without Global Spacing Activity

We found that Isw2 acts on specific targets through these specific transcription factor interactions, rather than acting on all nucleosomes genome-wide. We therefore speculate that Isw2 is in a globally repressed state in cells and activated solely on target nucleosomes. This inactivity is not consistent with work *in vitro*, and may be caused by a regulatory interaction that has not been previously observed in biochemical systems. For example, an unknown inhibitory factor that interacts with Isw2 or the nucleosome in cells may be lost during protein purification, allowing for the ubiquitous Isw2 chromatin remodeling activity observed *in vitro*. Additionally, it is conceivable that Isw2 is unable to bind to linker DNA the same way in a genomic context as it can bind *in vitro,* potentially due to the presence of unknown chromatin interacting components, molecular crowding, chromatin folding or other physiological differences not recapitulated *in vitro*. Maintaining Isw2 in an inactive state may allow organisms to conserve energy by controlling errant ATP hydrolysis while simultaneously enabling for rapid changes in chromatin structure and cellular output in differing contexts. It will be of great interest to determine how interactions with recruitment factor epitopes may alter the activity of Isw2 to elicit such precise nucleosome positioning outcomes in a cellular context.

### The Conserved WAC Domain Tethers Isw2 to Transcription Factor Epitopes in S. cerevisiae

The WAC domain, a broad N-terminal region of the Itc1 accessory protein, has been previously characterized as a DNA binding element that shares sequence conservation from flies to humans^49,52^. In this work, we have identified a previously undefined function of the WAC domain in mediating protein-protein interactions between ChRPs and transcription factors *in vivo.* We have further demonstrated that this mediation requires two conserved acidic residues within the WAC domain, which may allow future work to distinguish the DNA and protein binding capacity of the broader WAC domain region. Intriguingly, these critical acidic residues that are conserved between yeast, humans, mice, and fish have undergone an evolutionary charge reversal in *Drosophila*. It is conceivable that this charge reversal establishes a more general role in generating chromatin structure for the single ISWI-type protein found in flies, as opposed to the more specialized and context-dependent roles of the many ISWI-type ChRPs found in other organisms.

A recent model suggested strong interplay between the human Acf1 (yeast Itc1) N-terminus, extranucleosomal DNA and the histone H4 tail^50^. In this model, the human Acf1 N-terminus binds to extranucleosomal DNA in nucleosomes with long linker length, allowing the Snf2h (Isw2) catalytic subunit to engage the H4 tail. Snf2h engagement of the H4 tail relieves known autoinhibitory interactions^46,48^ thereby activating the remodeling complex. When linker DNA length shortens, the N-terminus of Acf1 switches to binding the H4 tail, thus displacing the Snf2h catalytic subunit and inactivating the complex through autoinhibition. This model was used to describe how ISWI complexes can be allosterically inactivated when linker DNA length shortens on nucleosomes and is a mechanistic model for how nucleosome length sensing can be achieved.

Our results indicate that the N-terminus of Itc1 does not have a primary cellular function of length sensing and nucleosome spacing. If the Itc1 N-terminus can bind H4 tail and transcription factor epitopes similarly to extranucleosomal DNA and H4 tail, the Hwang *et al* model^50^ can mechanistically explain the precise distance measurements made at targeted sites in cells. In this speculative model, Itc1 binds a targeting epitope at a genomic locus when the upstream nucleosome is far away. This orients the catalytic subunit on the appropriate nucleosome, which is remodeled toward the recruitment site. When the length between the nucleosome and the recruiting epitope is short enough, the Itc1 N-terminus may bind the H4 tail to inactivate Isw2 through autoinhibition. This function of binding the positively-charged H4 tail may be facilitated by the clustered acidic residues in the WAC domain, or may be mediated by another domain within the broadly-defined 374 base pair Itc1 N-terminus implicated in putative H4 tail binding. Determining whether this interplay between a transcription factor epitope and the H4 tail can tune distance measurements in cells will be important in future biophysical characterizations.

### The Benefits of Epitope-Mediated Chromatin Remodeling and an Interacting Barrier

What advantage may an interacting barrier provide that a general barrier cannot, particularly since a non-interacting barrier can still phase nucleosome arrays? We envision at least two major advantages of the interacting barrier model. First, an interacting barrier can behave directionally while a non-interacting barrier cannot. Indeed, Isw2 seems to be positioned on a specific barrier-proximal nucleosome through interactions between the WAC domain of Itc1 and the epitope on Ume6 or Swi6. Directionality allows for more refined establishment of transcriptionally relevant chromatin arrays. Second, an interacting barrier can be modulated in condition-specific manner through post-translational modification of the small epitope on the Isw2 recruitment factor. For example, one of the proteins that we identified as containing an Isw2-recruitment helix in yeast is Swi6, a critical regulator of the cell cycle in the G1/S transition. Interestingly, only three Swi6-regulated genes were identified as Swi6-mediated Isw2 recruitment sites. It is thus likely that the Swi6-Isw2 interaction can be tuned by cellular context, which is not possible for non-interacting barriers. Importantly, a tunable interacting barrier allows for continuous expression of the barrier and the ability to alter its barrier activity. This is a versatile mechanism through which chromatin structure may be spatiotemporally regulated in a dynamic fashion through these ChRP-recruitment factor interactions.

## Materials and Methods

### Yeast Strains and Plasmids

All yeast strains were derived from the parent strain *S. cerevisiae* W303 RAD5+. Gene deletions were made by replacing the gene of interest with antibiotic resistance markers amplified from pAG vectors. C-terminal deletions of genes were also made by replacing the region to be deleted with antibiotic resistance markers. N-terminal gene deletions were made by first replacing the region to be deleted with a URA3 marker, and then counterselecting with FOA to delete the URA3. Ume6-helix was introduced to yeast through plasmid transformation of a p416 vector containing the Ume6 helix fused to the SpyCatcher protein^41^. To make SpyTagged yeast strains, a C-terminal 3x-FLAG tag followed by the SpyTag sequence (AHIVMVDAYKPTK)^41^ was cloned into a pFA6a vector. Tags were then inserted at the endogenous locus of interest by homologous recombination of PCR products from the respective tagging vectors using selectable drug markers.

### Growth Conditions

Cells were grown at 30°C and 160 rpm in YPD (yeast extract-peptone-2% glucose) medium unless otherwise indicated. Strains were streaked from glycerol stocks onto 2% agar YPD plates and grown at 30°C for 2-3 days. An isolated colony was then grown overnight in 25 mL of YPD. This preculture was used to inoculate 25 mL of YPD at an OD_600_ of 0.2, which was grown to an OD_600_ of 0.6-0.8 for chromatin analysis. Yeast containing nonintegrating plasmids (p416) were grown in SD (-)Ura overnight, diluted to OD_600_=0.2 in YPD and grown to OD_600_=0.6-0.8 for chromatin analysis. Cells were then fixed with 1% formaldehyde and harvested for chromatin analysis.

### Protein Purification

Yeast strains containing the Isw2 variants of interest appended with a FLAG tag were grown at 30°C to an OD_600_ of ~1. Yeast were pelleted, washed with binding buffer (25mM HEPES pH 7.8, 300mM NaCl, 0.5mM EGTA, 0.1mM EDTA, 2mM MgCl_2_, 20% glycerol, 0.02% NP-40, 2mM betamercaptoethanol, 1mM PMSF, 1X protease inhibitor cocktail [Expedeon]) and then lysed via cryogrinding. Yeast powder was incubated with binding buffer for 90 minutes before the addition of 200ml bed volume anti-FLAG magnetic beads (Sigma M2). After 3 hour incubation at 4°C, beads were collected with magnets and washed 3 times with binding buffer and 3 times with elution buffer (25mM HEPES pH 7.8, 500mM NaCl, 0.5mM EGTA, 0.1mM EDTA, 2mM MgCl_2_, 20% glycerol, 0.02% NP-40, 2mM betamercaptoethanol, 1mM PMSF, 1X protease inhibitor cocktail [Expedeon]). A 0.5mg/ml solution of FLAG peptide in 100ml elution buffer was then added to the beads and allowed to incubate for 30 minutes. This process was repeated 3 more times for a total of 4 elutions. Elutions were analyzed by silver-staining and combined by estimated purity for aliquoting and storage at −80 degrees.

### Nucleosome Sliding Assay

Sliding assays were performed at least 3 independent times with reproducible results.

Recombinant yeast histones were purified as previously described^53^ and dialyzed by gradient salt dialysis onto the Widom 601 positioning sequence to create end-positioned nucleosomes with 60 base pairs of linker DNA^54^. Nucleosome sliding was performed at 25°C in sliding buffer (50mM KCl, 15mM HEPES, pH 7.8, 10mM MgCl_2_, 0.1mM EDTA, 5% sucrose, 0.2mg/ml bovine serum albumin [BSA], with or without 5mM ATP) by incubating 1ml or 1.5ml of purified Isw2 with 12.5nM reconstituted mononucleosomes for 40 minutes in 6ml reaction volume. Reactions were quenched by diluting 1:2 with solution containing 3mM competitor DNA and 5% sucrose. Native PAGE (6%) was used to separate the positioning of the mononucleosomes, with Cy5.5-labeled nucleosomal DNA detected by a LiCor Odyssey FC imager.

### Micrococcal Nuclease Digestions and Library Construction

Micrococcal nuclease digestions were performed with a minimum of two biological replicates as previously described^55^. Briefly, cells were grown to mid-log phase and fixed with 1% formaldehyde. Chromatin was digested with 10, 20, and 40 units of MNase for 10 minutes. Proper nuclease digestion of DNA was analyzed by agarose gel and samples with approximately 80% mononucleosomes were selected for library construction. After crosslink reversal, RNase treatment, Calf Intestine Phosphatase (CIP, NEB) treatment and Proteinase K digestion, mononucleosome-sized fragments were gel-purified and resulting DNA was used to construct libraries with the NuGEN Ovation Ultralow kit per the manufacturer’s instructions. Libraries were sequenced at the University of Oregon’s Genomics and Cell Characterization Core Facility on an Illumina NextSeq500 on the 37 cycle, paired-end, High Output setting, yielding approximately 10-20 million paired reads per sample.

### Chromatin Immunoprecipitation and Library Construction

Chromatin immunoprecipitation was performed with biological replicates as previously described^55^. Briefly, cells were grown to mid-log phase, fixed with 1% formaldehyde, and lysed by bead-beating in the presence of protease inhibitors. Chromatin was fragmented by shearing in a Bioruptor sonicator (Diagenode) for a total of 30 minutes (high output, 3×10’ cycles of 30 sec. on, 30 sec. off). Sonication conditions were optimized to produce an average fragment size of ∼300 basepairs. FLAG-tagged protein was immunoprecipitated using FLAG antibody (Sigma) and Protein G magnetic beads (Invitrogen). After crosslink reversal and Proteinase K digestion, DNA was purified using Qiagen MinElute columns and quantified by Qubit High-Sensitivity fluorometric assay. Libraries were prepared using the NuGEN Ovation Ultralow kit by the manufacturer’s instructions and sequenced at the University of Oregon’s Genomics and Cell Characterization Core Facility on an Illumina NextSeq500 with 37 cycles of paired-end setting, yielding approximately 10 million single-end reads per sample. Only the first read (R1) of each paired read was taken for downstream alignments and processing.

### RNA Extraction and Library Construction

For RNA-Seq (minimum two biological replicates), RNA was purified by hot acid phenol extraction followed by polyA selection and strand-specific library construction using the NuGEN Universal Plus mRNA Kit according to the manufacturer’s instructions. Libraries were sequenced on an Illumina NextSeq500 on the 37 cycle, paired-end, High Output setting. Paired end reads were quality filtered for adapter contamination and low quality ends using trimmomatic^56^. After quality filtering an average of 10.5 million reads per paired end sample remained. Surviving reads were mapped to the *S. cerevisiae* reference genome^57^ using STAR (V.2.5.3)^58^. Gene counts were quantified from uniquely aligning reads using HTSeq (V.0.9.1)^59^. Differential gene expression was performed using DESeq2(V.1.22.2)^60^, and expression graphs were generated using ggplot2^61^.

### Data Processing and Analysis

MNase sequencing data were analyzed as described previously^62^. Briefly, paired-end reads were aligned to the *S. cerevisiae* reference genome^57^ with Bowtie 2^63^, and filtered computationally for unique fragments between 100 and 200 bp. Dyad positions were calculated as the midpoint of paired reads, then dyad coverage was normalized across the *S. cerevisiae* genome for an average read/bp of 1.0. Dyad coverage is displayed in all figures. Nucleosome alignments to transcription Ume6 binding sites were performed by taking average dyad signal at each position relative to all 202 intergenic instances of a Ume6 motif center *(WNGGCGGCWW).* Preinitiation complex (PIC) locations were obtained from Rhee et. al.^64^. For ChIP-Seq data, single-end reads were aligned to the *S. cerevisiae* reference genome with Bowtie 2 and total read coverage was normalized such that the average read at a genomic location was 1.0. ChIP peaks were called using a 400 bp sliding window with a threshold average enrichment within the window of 3.0. Data were visualized using Integrated Genome Browser^65^. The datasets generated during this study are available in the GEO Database with accession code GSE149804.

## Author Contributions and Notes

Conceptualization, D.A.D. and J.N.M.; Methodology, D.A.D., J.G.C., V.N.T., A.L.V., T.B.B., L.E.M., and J.N.M.; Investigation, D.A.D., J.G.C., V.N.T., A.L.V., T.B.B., L.E.M., and J.N.M.; Writing – Original Draft, D.A.D. and J.N.M.; Writing – Reviewing & Editing, D.A.D., J.G.C., L.E.M., and J.N.M.; Visualization, D.D. and J.N.M.; Supervision, L.E.M. and J.N.M.; Project Administration, J.N.M.; Funding Acquisition, J.N.M.

The authors declare no conflict of interest.

Sequencing data sets can be accessed in the Gene Expression Omnibus with Accession Number GSE149804.

## Acknowledgments

The authors thank Christine Cucinotta for helpful comments on the manuscript, and Greg Bowman for helpful discussions about the project. This work was supported by NIH training grants T32 GM007759 (to D.A.D.) and T32 GM007413 (to D.A.D. and V.N.T.), and by NIGMS grant R01 GM129242 (J.N.M.), the Donald and Delia Baxter Foundation (J.N.M.) and the Medical Research Foundation of Oregon (J.N.M.).

---

**Supplemental Figure S1 (related to Figure 1).**
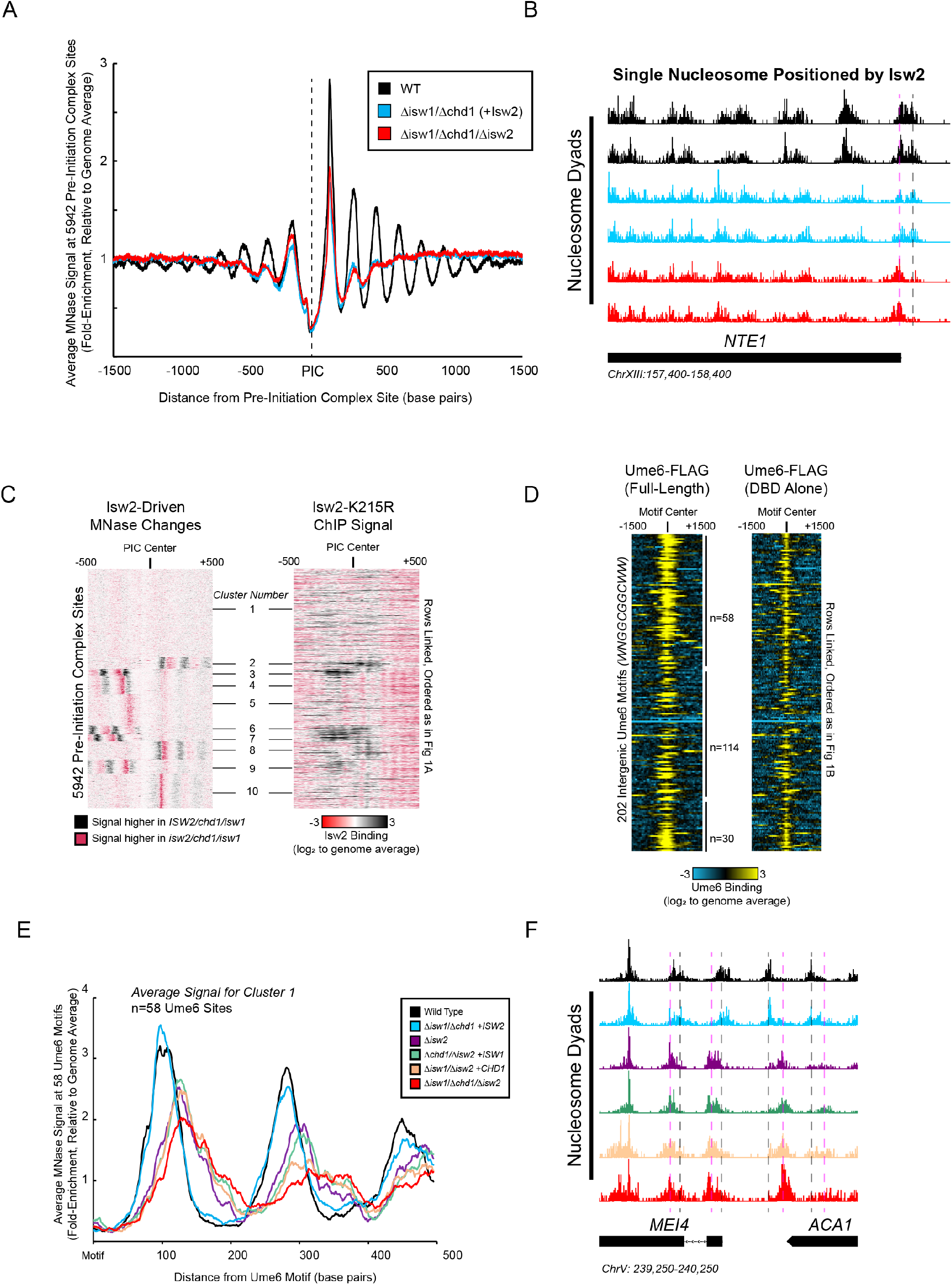
Isw2 is a Precise Specialist at Target Nucleosomes. **A** Meta-analysis nucleosome dyad signal at all 5942 PIC sites shows that Isw2 confers no global nucleosome organizing activity throughout yeast cells. **B** Example Isw2 target locus where addition of Isw2 in the absence of Isw1 and Chd1 leads to positioning of a single motif-proximal nucleosome at the *NTE1* locus. Vertical gray dashed lines indicate the wild-type nucleosome locations while vertical dashed pink lines indicate an alternate *isw2* deletion strain position at the motif-proximal nucleosome. The *ISW2/chd1/isw1* strain can position the motif proximal nucleosome, but all distal nucleosomes are disorganized much like the *isw2/chd1/isw1* strain. **C** Heat map showing that PIC clusters from Figure 1A that display Isw2-dependent nucleosome changes overlap with regions where Isw2(K215R)-FLAG ChIP signal is present. All rows are linked and ordered identically to Figure 1A. **D** Heat maps showing that full length Ume6-FLAG and Ume6(Δ2-763)-FLAG associate with similar targets. All rows are linked and all 202 intergenic Ume6 motifs are displayed. **E** Metaanalysis of nucleosome dyad signal for indicated strains at the 58 Ume6 target loci associated with cluster 1 in Figure 1B demonstrates that Isw2 but not Chd1 or Isw1 is necessary and sufficient to position nucleosomes at Ume6 target loci. **F** Genome Browser image showing nucleosome dyad signal for indicated strains at the MEI4-ACA1 locus, a representative Ume6 target locus. The color schemes are shared between Figures S1E and S1F according to the key between figures. Vertical gray lines indicate wild-type nucleosome positions, while vertical pink lines indicate ectopic nucleosome positions associated with loss of Isw2 activity.

**Supplemental Figure S2 (related to Figure 2).**
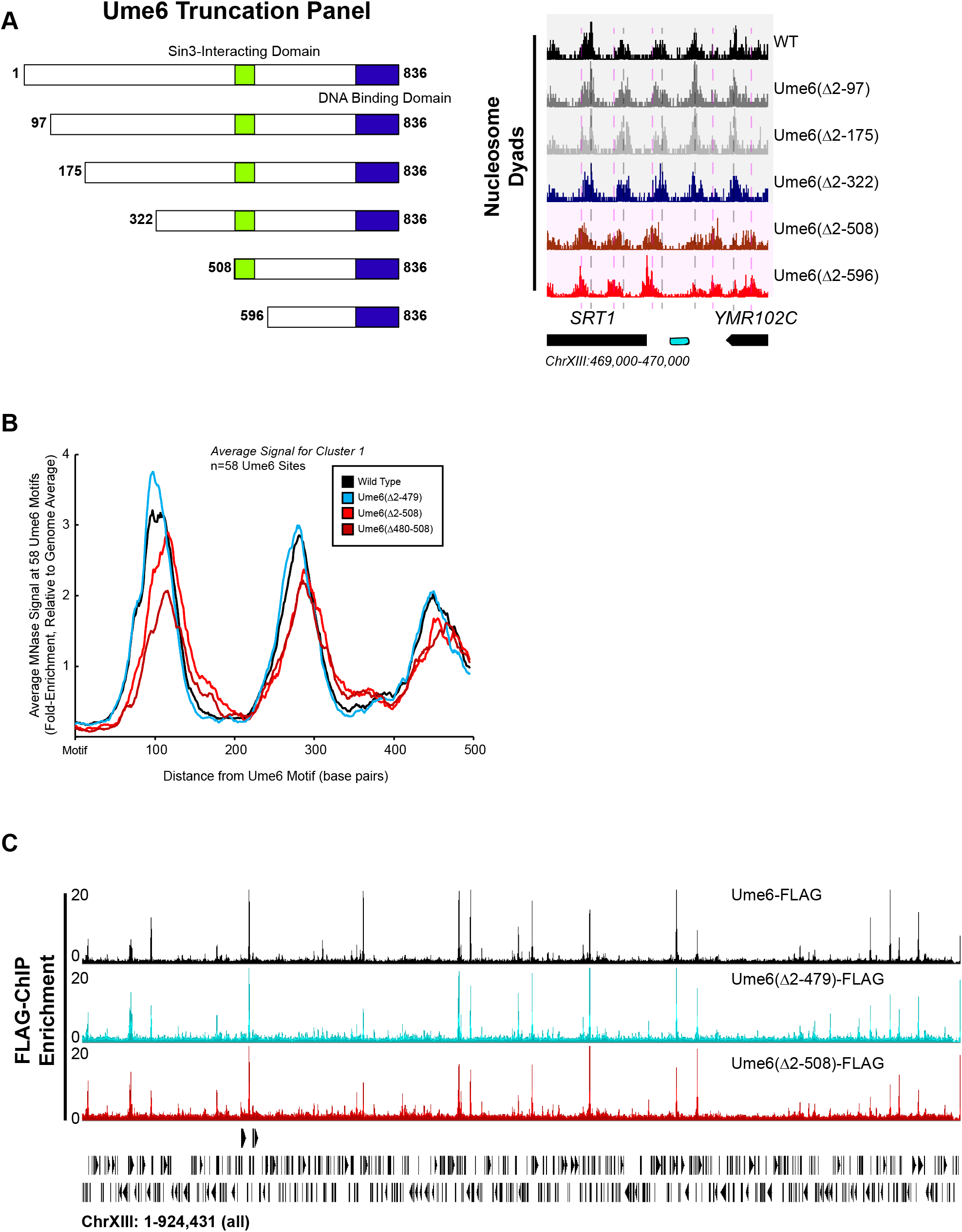
The Ume6 Helix Between Residues 479 and 508 Recruits Isw2 to Ume6 Targets. **A** (left) Schematic representation of the truncation panel initially used to identify the region of Ume6 required for Isw2 recruitment. Green square indicates the part of Ume6 known to recruit SIN3, while the blue rectangle indicates the Ume6 DNA binding domain. (right) Genome Browser image for a representative locus showing nucleosome dyad signal for the indicated strains. Vertical gray lines denote wild-type nucleosome positions while vertical pink lines indicate ectopic nucleosome positions associated with truncations beyond 322 N-terminal amino acids. The Isw2-recruitment domain was thus determined to be between residues 322 and 508 in Ume6. **B** Meta-analysis of nucleosome dyad signal at the 58 Ume6 target loci associated with cluster 1 in Figure 1B showing loss of residues 479-508 from Ume6 results in ectopic nucleosome positioning while N-terminal deletion of residues 2-479 preserves wild-type nucleosome positioning at Ume6 sites. **C** Genome Browser image demonstrating no loss in Ume6-ChIP signal for relevant Ume6 truncations across all of chromosome XIII. Signal is read-corrected enrichment, relative to genome average.

**Supplemental Figure S3 (related to Figure 2).**
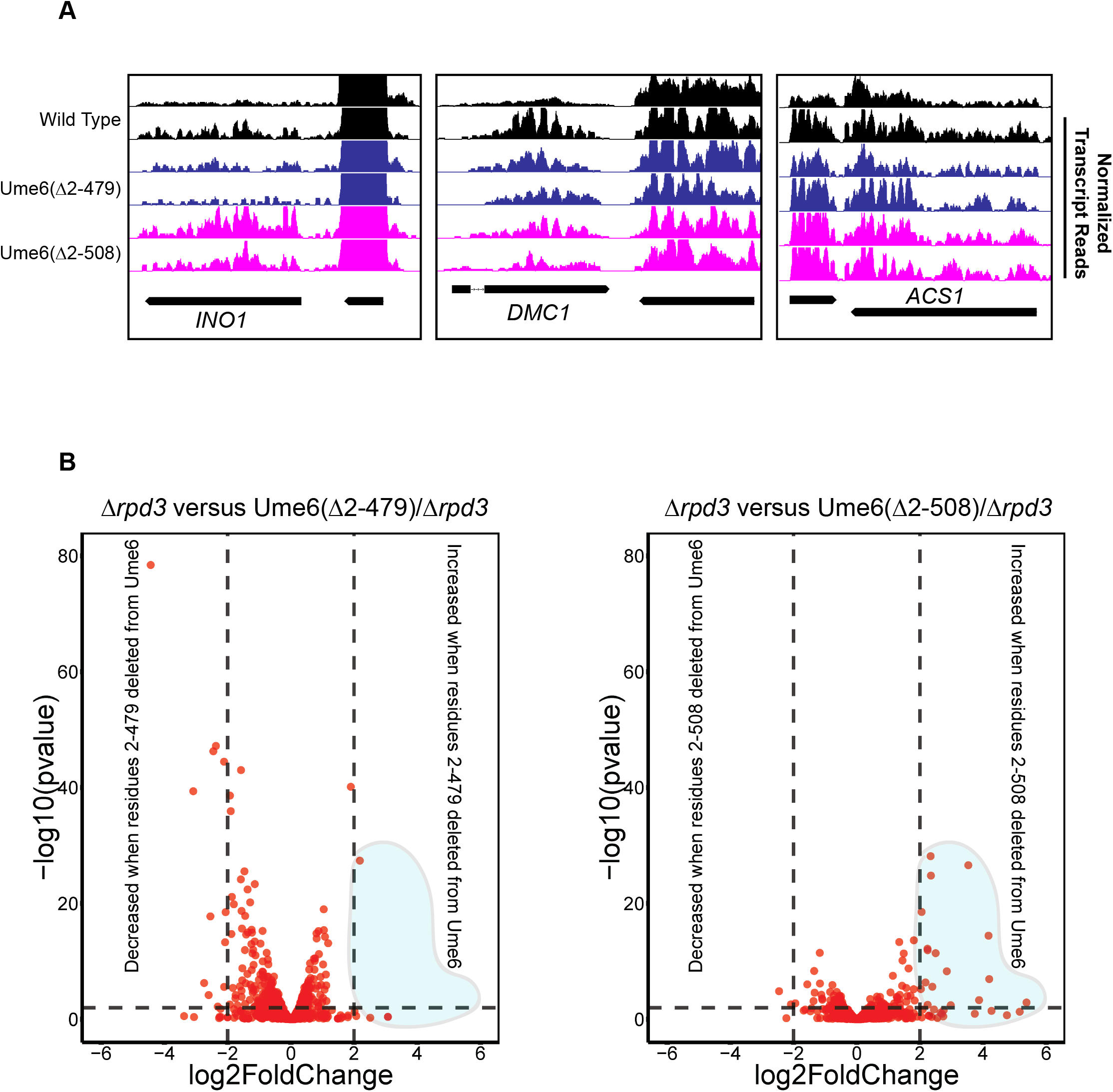
Transcription Data Support a Role of Ume6 Residues 479-508 for Isw2 Recruitment and not Rpd3 Activity. **A** Genome Browser images showing transcript abundance for the three representative Ume6 target loci for indicated Ume6 truncation strains in the presence of functional Rpd3, demonstrating the expected very subtle increase in transcription associated with loss of Isw2 at the INO1 locus. Biological replicates for each strain are shown. **B** Volcano plots for genes containing Ume6 motifs showing log2 change in transcription and associated statistical significance for indicated strains. Horizontal dashed line indicates a p-value of 0.01. Vertical dashed lines indicate a change in transcription of +/− 4-fold. Retention of residues 479-508 prevents large-scale increases in transcription associated with deletion of Rpd3 and Isw2 (see blue shaded region).

**Supplemental Figure S4 (related to Figure 2).**
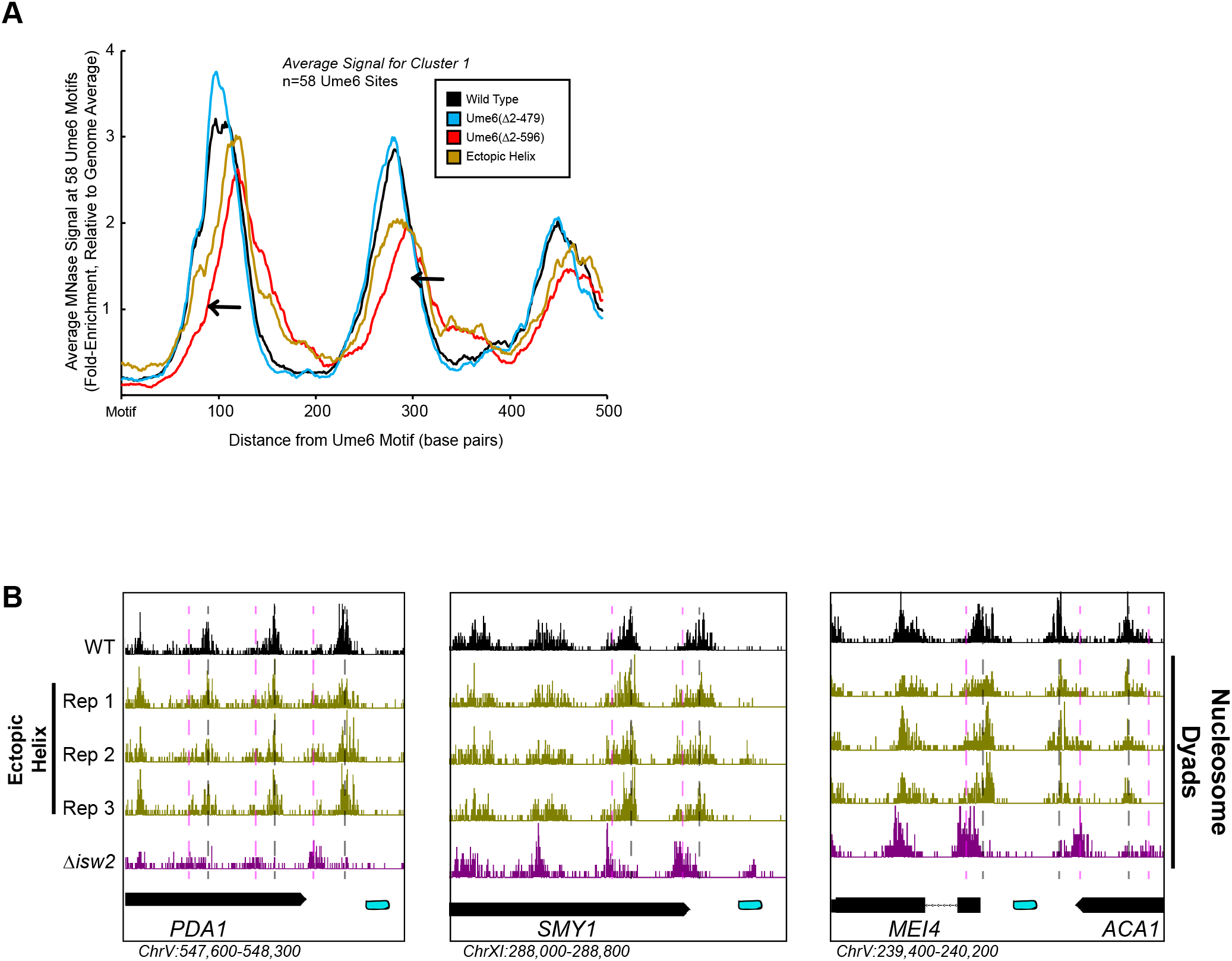
Ectopic Display of the Ume6 Helical Element can Rescue Isw2 Activity at Ume6 Targets. **A** Meta-analysis of nucleosome dyad signal at the 58 Ume6 target loci associated with cluster 1 in Figure 1B showing partial rescue of nucleosome positioning when residues 479-508 are ectopically displayed on the C-terminus of Ume6 Δ2-596 through SpyTag-SpyCatcher pairs. Black arrows indicate wild-type signal gained by ectopic display of the recruitment helix. **B** Genome Browser image showing additional replicates where SpyTag/SpyCatcher-mediated ectopic tethering of Ume6 residues 479-508 recovers wild-type nucleosome positions at Ume6 targets (as in Figure 2C).

**Supplemental Figure S5 (related to Figure 3).**
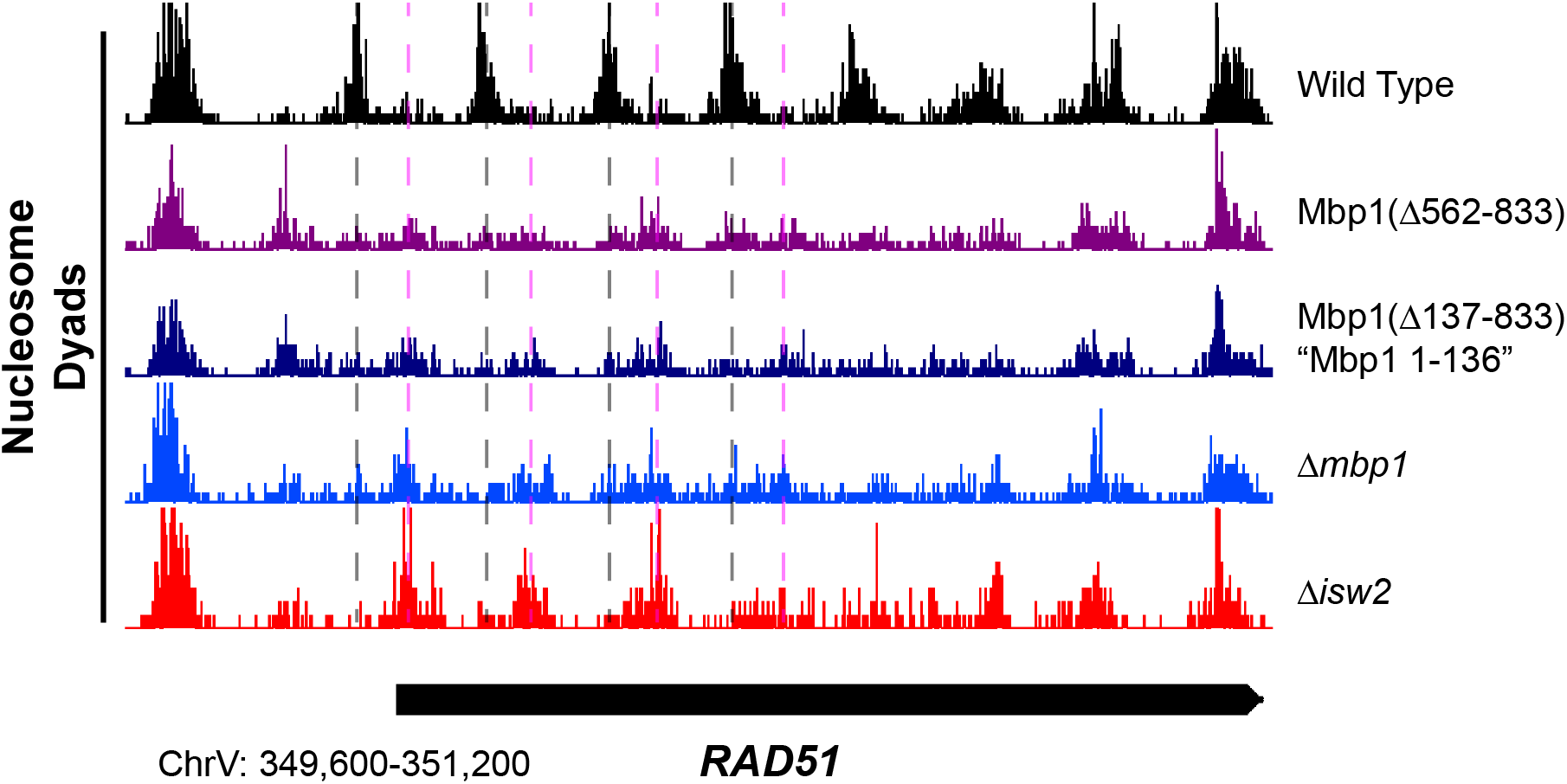
Truncation of the Mbp1 C-terminus Eliminates Isw2-Directed Nucleosome Positioning at Mbp1 Targets. Genome Browser image showing nucleosome positioning at the *RAD51* locus for indicated strains. Dashed vertical gray lines denote positions of nucleosomes in the wild-type strain, while dashed vertical pink lines show positions of nucleosomes in the absence of functional Isw2.

**Supplemental Figure S6 (related to Figure 5).**
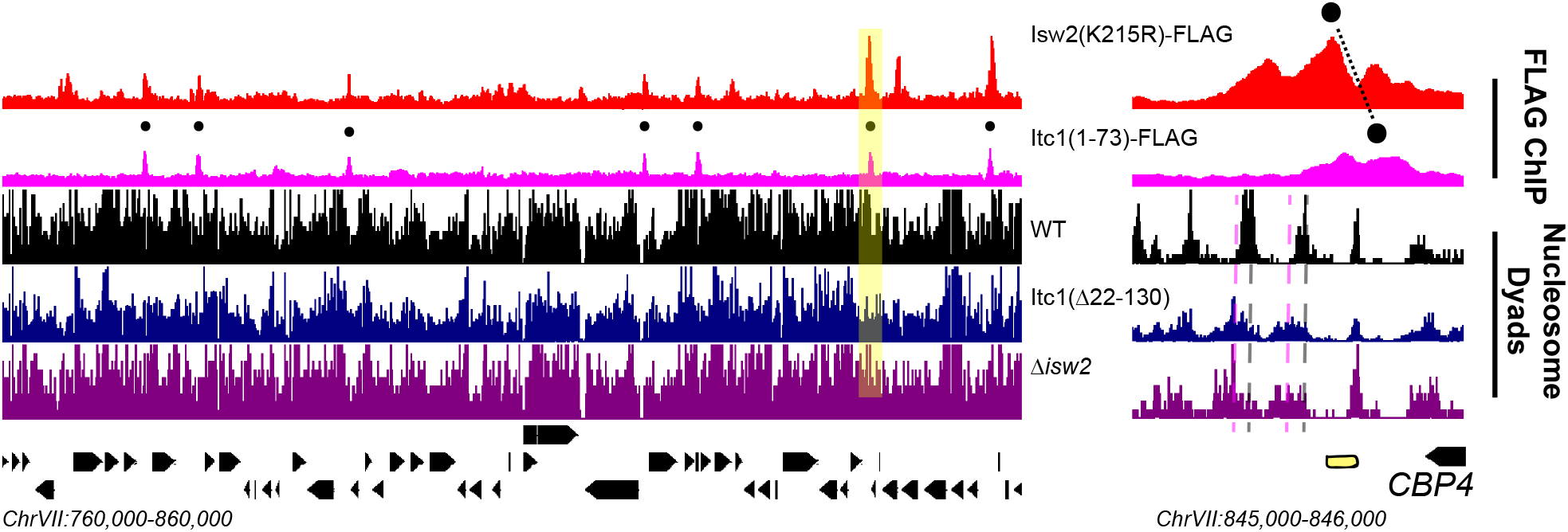
The WAC Domain Orients the Isw2 Catalytic Domain at Nearly Half of Detected Isw2 Targets in Yeast. (left) Genome Browser image showing Isw2(K215R)-FLAG and Itc1(1-73)-FLAG ChIP signal and wild type, Itc1(Δ22-130), and Δ*isw2* nucleosome dyad signal for a 100kb section of chromosome VII. Black circles indicate genomic loci where Isw2(K215R) and Itc1(1-73) ChIP signal overlap. (right) Zoomed Genome Browser image of the section highlighted in yellow showing offset nature of the Isw2 peak and Itc1 peak. Black circles connected by a black line indicates the offset nature of the Isw2 and Itc1 ChIP peaks. The shifted nucleosome is to the left of the Itc1-Isw2 axis.

## References

1. Luger, K., Mader, A.W., Richmond, R.K., Sargent, D.F. & Richmond, T.J. Crystal structure of the nucleosome core particle at 2.8 A resolution. Nature 389, 251–60 (1997).

2. Kornberg, R.D. Chromatin structure: a repeating unit of histones and DNA. Science 184, 868–71 (1974).

3. Lai, W.K.M. & Pugh, B.F. Understanding nucleosome dynamics and their links to gene expression and DNA replication. Nat Rev Mol Cell Biol 18, 548–562 (2017).

4. Zhou, C.Y., Johnson, S.L., Gamarra, N.I. & Narlikar, G.J. Mechanisms of ATP-Dependent Chromatin Remodeling Motors. Annu Rev Biophys 45, 153–81 (2016).

5. Clapier, C.R., Iwasa, J., Cairns, B.R. & Peterson, C.L. Mechanisms of action and regulation of ATP-dependent chromatin-remodelling complexes. Nat Rev Mol Cell Biol 18, 407–422 (2017).

6. Stockdale, C., Flaus, A., Ferreira, H. & Owen-Hughes, T. Analysis of nucleosome repositioning by yeast ISWI and Chd1 chromatin remodeling complexes. J Biol Chem 281, 16279–88 (2006).

7. Hauk, G., McKnight, J.N., Nodelman, I.M. & Bowman, G.D. The chromodomains of the Chd1 chromatin remodeler regulate DNA access to the ATPase motor. Mol Cell 39, 711–23 (2010).

8. McKnight, J.N., Jenkins, K.R., Nodelman, I.M., Escobar, T. & Bowman, G.D. Extranucleosomal DNA binding directs nucleosome sliding by Chd1. Mol Cell Biol 31, 4746–59 (2011).

9. Kagalwala, M.N., Glaus, B.J., Dang, W., Zofall, M. & Bartholomew, B. Topography of the ISW2-nucleosome complex: insights into nucleosome spacing and chromatin remodeling. EMBO J 23, 2092–104 (2004).

10. Lusser, A., Urwin, D.L. & Kadonaga, J.T. Distinct activities of CHD1 and ACF in ATP-dependent chromatin assembly. Nat Struct Mol Biol 12, 160–6 (2005).

11. Tsukiyama, T., Palmer, J., Landel, C.C., Shiloach, J. & Wu, C. Characterization of the imitation switch subfamily of ATP-dependent chromatin-remodeling factors in Saccharomyces cerevisiae. Genes Dev 13, 686–97 (1999).

12. Pointner, J. et al. CHD1 remodelers regulate nucleosome spacing in vitro and align nucleosomal arrays over gene coding regions in S. pombe. EMBO J 31, 4388–403 (2012).

13. Lee, W. et al. A high-resolution atlas of nucleosome occupancy in yeast. Nat Genet 39, 1235–44 (2007).

14. Mavrich, T.N. et al. Nucleosome organization in the Drosophila genome. Nature 453, 358–62 (2008).

15. Valouev, A. et al. Determinants of nucleosome organization in primary human cells. Nature 474, 516–20 (2011).

16. Krietenstein, N. et al. Genomic Nucleosome Organization Reconstituted with Pure Proteins. Cell 167, 709–721 e12 (2016).

17. Wiechens, N. et al. The Chromatin Remodelling Enzymes SNF2H and SNF2L Position Nucleosomes adjacent to CTCF and Other Transcription Factors. PLoS Genet 12, e1005940 (2016).

18. Baldi, S. et al. Genome-wide Rules of Nucleosome Phasing in Drosophila. Mol Cell 72, 661–672 e4 (2018).

19. Gkikopoulos, T. et al. A role for Snf2-related nucleosome spacing enzymes in genome-wide nucleosome organization. Science 333, 1758–60 (2011).

20. Zhang, Z. et al. A packing mechanism for nucleosome organization reconstituted across a eukaryotic genome. Science 332, 977–80 (2011).

21. Mavrich, T.N. et al. A barrier nucleosome model for statistical positioning of nucleosomes throughout the yeast genome. Genome Res 18, 1073–83 (2008).

22. Yan, C., Chen, H. & Bai, L. Systematic Study of Nucleosome-Displacing Factors in Budding Yeast. Mol Cell 71, 294–305 e4 (2018).

23. Gelbart, M.E., Bachman, N., Delrow, J., Boeke, J.D. & Tsukiyama, T. Genome-wide identification of Isw2 chromatin-remodeling targets by localization of a catalytically inactive mutant. Genes Dev 19, 942–54 (2005).

24. Goldmark, J.P., Fazzio, T.G., Estep, P.W., Church, G.M. & Tsukiyama, T. The Isw2 chromatin remodeling complex represses early meiotic genes upon recruitment by Ume6p. Cell 103, 423–33 (2000).

25. Fazzio, T.G. et al. Widespread collaboration of Isw2 and Sin3-Rpd3 chromatin remodeling complexes in transcriptional repression. Mol Cell Biol 21, 6450–60 (2001).

26. Yadon, A.N., Singh, B.N., Hampsey, M. & Tsukiyama, T. DNA looping facilitates targeting of a chromatin remodeling enzyme. Mol Cell 50, 93–103 (2013).

27. Donovan, D.A. et al. Engineered Chromatin Remodeling Proteins for Precise Nucleosome Positioning. Cell Rep 29, 2520–2535 e4 (2019).

28. Bowman, G.D. & McKnight, J.N. Sequence-specific targeting of chromatin remodelers organizes precisely positioned nucleosomes throughout the genome. Bioessays 39, 1–8 (2017).

29. McKnight, J.N., Tsukiyama, T. & Bowman, G.D. Sequence-targeted nucleosome sliding in vivo by a hybrid Chd1 chromatin remodeler. Genome Res 26, 693–704 (2016).

30. Dang, W. & Bartholomew, B. Domain architecture of the catalytic subunit in the ISW2-nucleosome complex. Mol Cell Biol 27, 8306–17 (2007).

31. Dang, W., Kagalwala, M.N. & Bartholomew, B. Regulation of ISW2 by concerted action of histone H4 tail and extranucleosomal DNA. Mol Cell Biol 26, 7388–96 (2006).

32. Hota, S.K. et al. Nucleosome mobilization by ISW2 requires the concerted action of the ATPase and SLIDE domains. Nat Struct Mol Biol 20, 222–9 (2013).

33. Kassabov, S.R., Henry, N.M., Zofall, M., Tsukiyama, T. & Bartholomew, B. High-resolution mapping of changes in histone-DNA contacts of nucleosomes remodeled by ISW2. Mol Cell Biol 22, 7524–34 (2002).

34. Zofall, M., Persinger, J. & Bartholomew, B. Functional role of extranucleosomal DNA and the entry site of the nucleosome in chromatin remodeling by ISW2. Mol Cell Biol 24, 10047–57 (2004).

35. Zofall, M., Persinger, J., Kassabov, S.R. & Bartholomew, B. Chromatin remodeling by ISW2 and SWI/SNF requires DNA translocation inside the nucleosome. Nat Struct Mol Biol 13, 339–46 (2006).

36. Ocampo, J., Chereji, R.V., Eriksson, P.R. & Clark, D.J. The ISW1 and CHD1 ATP-dependent chromatin remodelers compete to set nucleosome spacing in vivo. Nucleic Acids Res 44, 4625–35 (2016).

37. Li, M. et al. Dynamic regulation of transcription factors by nucleosome remodeling. Elife 4(2015).

38. Kubik, S. et al. Opposing chromatin remodelers control transcription initiation frequency and start site selection. Nat Struct Mol Biol 26, 744–754 (2019).

39. Kelley, L.A., Mezulis, S., Yates, C.M., Wass, M.N. & Sternberg, M.J. The Phyre2 web portal for protein modeling, prediction and analysis. Nat Protoc 10, 845–58 (2015).

40. Washburn, B.K. & Esposito, R.E. Identification of the Sin3-binding site in Ume6 defines a two-step process for conversion of Ume6 from a transcriptional repressor to an activator in yeast. Mol Cell Biol 21, 2057–69 (2001).

41. Zakeri, B. et al. Peptide tag forming a rapid covalent bond to a protein, through engineering a bacterial adhesin. Proc Natl Acad Sci U S A 109, E690–7 (2012).

42. Koch, C., Moll, T., Neuberg, M., Ahorn, H. & Nasmyth, K. A role for the transcription factors Mbp1 and Swi4 in progression from G1 to S phase. Science 261, 1551–7 (1993).

43. Breeden, L. Start-specific transcription in yeast. Curr Top Microbiol Immunol 208, 95–127 (1996).

44. Nair, M. et al. NMR structure of the DNA-binding domain of the cell cycle protein Mbp1 from Saccharomyces cerevisiae. Biochemistry 42, 1266–73 (2003).

45. Foord, R., Taylor, I.A., Sedgwick, S.G. & Smerdon, S.J. X-ray structural analysis of the yeast cell cycle regulator Swi6 reveals variations of the ankyrin fold and has implications for Swi6 function. Nat Struct Biol 6, 157–65 (1999).

46. Clapier, C.R. & Cairns, B.R. Regulation of ISWI involves inhibitory modules antagonized by nucleosomal epitopes. Nature 492, 280–4 (2012).

47. Yan, L., Wang, L., Tian, Y., Xia, X. & Chen, Z. Structure and regulation of the chromatin remodeller ISWI. Nature 540, 466–469 (2016).

48. Ludwigsen, J. et al. Concerted regulation of ISWI by an autoinhibitory domain and the H4 N-terminal tail. Elife 6(2017).

49. Fyodorov, D.V. & Kadonaga, J.T. Binding of Acf1 to DNA involves a WAC motif and is important for ACF-mediated chromatin assembly. Mol Cell Biol 22, 6344–53 (2002).

50. Hwang, W.L., Deindl, S., Harada, B.T. & Zhuang, X. Histone H4 tail mediates allosteric regulation of nucleosome remodelling by linker DNA. Nature 512, 213–7 (2014).

51. Yen, K., Vinayachandran, V., Batta, K., Koerber, R.T. & Pugh, B.F. Genome-wide nucleosome specificity and directionality of chromatin remodelers. Cell 149, 1461–73 (2012).

52. Ito, T. et al. ACF consists of two subunits, Acf1 and ISWI, that function cooperatively in the ATP-dependent catalysis of chromatin assembly. Genes Dev 13, 1529–39 (1999).

53. Luger, K., Rechsteiner, T.J. & Richmond, T.J. Expression and purification of recombinant histones and nucleosome reconstitution. Methods Mol Biol 119, 1–16 (1999).

54. Lowary, P.T. & Widom, J. New DNA sequence rules for high affinity binding to histone octamer and sequence-directed nucleosome positioning. J Mol Biol 276, 19–42 (1998).

55. Rodriguez, J., McKnight, J.N. & Tsukiyama, T. Genome Wide Analysis of Nucleosome Positions, Occupancy, and Accessibility in Yeast: Nucleosome Mapping, High-Resolution Histone ChIP, and NCAM. Curr Protoc Mol Biol 108, 21 28 1–16 (2014).

56. Bolger, A.M., Lohse, M. & Usadel, B. Trimmomatic: a flexible trimmer for Illumina sequence data. Bioinformatics 30, 2114–20 (2014).

57. Cunningham, F. et al. Ensembl 2015. Nucleic Acids Res 43, D662–9 (2015).

58. Dobin, A. et al. STAR: ultrafast universal RNA-seq aligner. Bioinformatics 29, 15–21 (2013).

59. Anders, S., Pyl, P.T. & Huber, W. HTSeq--a Python framework to work with high-throughput sequencing data. Bioinformatics 31, 166–9 (2015).

60. Love, M.I., Huber, W. & Anders, S. Moderated estimation of fold change and dispersion for RNA-seq data with DESeq2. Genome Biol 15, 550 (2014).

61. Wickham, H. ggplot2: Elegant Graphics for Data Analysis. Springer-Verlag (2016).

62. McKnight, J.N. & Tsukiyama, T. The conserved HDAC Rpd3 drives transcriptional quiescence in S. cerevisiae. Genom Data 6, 245–8 (2015).

63. Langmead, B. & Salzberg, S.L. Fast gapped-read alignment with Bowtie 2. Nat Methods 9, 357–9 (2012).

64. Rhee, H.S. & Pugh, B.F. Genome-wide structure and organization of eukaryotic pre-initiation complexes. Nature 483, 295–301 (2012).

65. Freese, N.H., Norris, D.C. & Loraine, A.E. Integrated genome browser: visual analytics platform for genomics. Bioinformatics 32, 2089–95 (2016).

